# Prenatal exposure to EDCs dis-integrates and reconstitutes neuromolecular-behavioral relationships in adult rats

**DOI:** 10.1101/2020.10.12.335984

**Authors:** Morgan E. Hernandez Scudder, Rebecca L. Young, Lindsay M. Thompson, Pragati Kore, David Crews, Hans A. Hofmann, Andrea C. Gore

## Abstract

Exposure to endocrine-disrupting chemicals (EDCs) is ubiquitous in all species, including humans. Previous studies have shown behavioral deficits caused by EDCs that have implications for social competence and sexual selection. The neuromolecular mechanisms for these behavioral changes induced by EDCs have not been thoroughly explored. Here, we tested the hypothesis that EDCs administered to rats during a critical period of embryonic brain development would lead to disruption of normal social preference behavior, and that this involves a network of underlying gene pathways in brain regions that regulate these behaviors. Rats were exposed prenatally to human-relevant concentrations of EDCs [polychlorinated biphenyls (PCB), an industrial chemical mixture; vinclozolin (VIN), a fungicide], or vehicle. In adulthood, a sociosexual preference test (choice between hormone-primed and hormone-depleted opposite-sex rats) was administered. We profiled gene expression of in three brain regions involved in these behaviors [preoptic area (POA), medial amygdala (MeA), ventromedial nucleus (VMN)]. Prenatal PCBs impaired sociosexual preference in both sexes, and VIN disrupted this behavior in males. Each brain region (POA, MeA, VMN) had unique sets of genes altered in a sex- and EDC-specific manner. Sexually dimorphic gene expression disruption was particularly prominent for gene modules pertaining to sex steroid hormones and nonapeptides in the MeA. EDC exposure also changed the relationships between gene expression and behavior in the mate preference test, a pattern we refer to as dis-integration and reconstitution. These findings underscore the profound effects that developmental exposure to EDCs can have on adult social behavior, highlight sex-specific and individual variation in responses, and provide a foundation for further work on the disruption of mate preference behavior after prenatal exposure to EDCs.

## Introduction

Environmental contamination with endocrine-disrupting chemicals (EDCs) perturbs hormones and their actions in virtually all species and ecosystems (Gore et al., 2015). Prenatal EDC exposures pose a particular risk due to the exquisite sensitivity of the developing brain to gonadal hormones, which are required for sex-typical differentiation and development of neural circuits, and the manifestation of behaviors. In the hypothalamus of male rodents and other mammals, prenatal and early postnatal testicular hormones masculinize and defeminize circuits. In females, the relative quiescence of the ovary and concomitantly lower gonadal hormone production, together with alpha-fetoprotein that prevents estrogens’ crossing the blood-brain-barrier, is responsible for brain feminization and demasculinization (Bakker et al., 2006; Nugent et al., 2015; Schwarz & McCarthy, 2008; Wright, Schwarz, Dean, & McCarthy, 2010).

The effects of developmental EDC exposure on sexually dimorphic social behaviors and gene expression patterns in different brain regions have been described for several classes of chemicals. Although individual EDCs are not pure hormone agonists or antagonists, some [such as certain polychlorinated biphenyls (PCBs) and bisphenol A (BPA)] mimic or disrupt estrogen signaling (Dickerson & Gore, 2007), and others [vinclozolin (VIN) and phthalates] are anti-androgenic (Euling et al., 2002; Stroheker et al., 2005). PCBs, widespread industrial chemical contaminants, alter gene expression in the hypothalamus (Dickerson, Cunningham, & Gore, 2011; Faass, Ceccatelli, Schlumpf, & Lichtensteiger, 2013; Topper et al., 2019) and change interactions of adult rats with conspecifics (Hernandez Scudder et al., 2020; Bell, Hart, & Gore, 2016; Colciago et al., 2009; Cummings, Clemens, & Nunez, 2008; Steinberg, Juenger, & Gore, 2007). Bisphenol A (BPA) from plastic, and the fungicide VIN also change brain gene expression (BPA: Wolstenholme et al., 2012, VIN: Skinner, Savenkova, Zhang, Gore, & Crews, 2014; Faass et al., 2013; Lichtensteiger et al., 2015) and sociosexual behavior (BPA: Jones, Shimell, & Watson, 2011; Monje, Varayoud, Muñoz-de-Toro, Luque, & Ramos, 2009; Porrini et al., 2005, VIN: Colbert et al., 2005; Krishnan et al., 2018). Prenatal exposure to phthalates causes long-lasting changes to gene expression in the hypothalamus and beyond (Gao et al., 2018; Lin et al., 2015). Phthalate exposure early in life also cause deficits in cognitive and social behaviors (Lin et al., 2015; R. Wang, Xu, & Zhu, 2016). In most cases, outcomes are dependent on the dose, timing, and length of exposure, as well as the sex of the animal. This is not surprising considering the dynamic nature of endogenous hormone signaling as the brain develops, and the vulnerability of estrogenic and androgenic pathways to EDCs.

Reproductive success is contingent upon sex-appropriate differentiation of the brain during early life. For an individual to reproduce successfully, appropriate dyadic interactions with another sexually mature potential mate of the opposite sex are required. This process involves assessment of an opposite-sex animal’s fitness through a variety of physical and behavioral cues, including hormonal status, as well-documented in rats (Drewett, 1973; Edwards & Einhorn, 1986; Eliasson & Meyerson, 1975). There is plasticity in this behavior, with the decision-making process affected by prior sexual experience of both individuals, estrous cycle stage, hormone levels, and other factors. Within the brain, a complex social decision-making network (O’Connell & Hofmann, 2012) comprising hypothalamic [e.g., ventromedial nucleus (VMN), preoptic area (POA)] and extra-hypothalamic [e.g., medial amygdala (MeA)] regions expresses specific genes and proteins that modulate these behaviors (Spiteri et al., 2010).

Here, we tested the hypothesis that prenatal EDC exposures would cause disruptions to the pattern of expression of a suite of genes in three brain regions in the social decision-making network (VMN, POA, MeA) and that this underlies functional deficits in an ethologically-relevant sociosexual behavioral task. Previous work has not considered the complex inter-relationships of these phenotypes, a gap we intended to fill in current work. The goal was to determine whether these relationships would break down (become “dis-organized”) and/or become reconstituted into novel patterns. To do this, we combined an integrative analysis of behavioral and hormonal phenotypes and gene co-expression patterns to characterize relationships between multiple measures of motivated behavior, gene expression patterns, and circulating hormone levels in response to prenatal EDC exposure in both male and female rats.

## Methods and materials

### Experimental design

All rat procedures were conducted in compliance with protocols approved by IACUC at The University of Texas at Austin. Sprague-Dawley rats purchased from Envigo (Houston) were housed in colony rooms with consistent temperature (22°C) and light cycle (14:10 dark:light, lights off at 1100). All rats had *ad libitum* access to water and were fed a low phytoestrogen rat chow (Teklad 2019, Envigo).

To generate experimental rats, virgin females were mated with sexually experienced males. Successful mating was indicated by the presence of sperm in a vaginal smear. The day after mating overnight was termed embryonic day 1 (E1). Pregnant rats received intraperitoneal (i.p.) injections of one of three treatments daily from E8-E18: (1) Vehicle (6% DMSO in sesame oil), (2) A1221 (1mg/kg), or (3) VIN (1mg/kg). Each dam was exposed to the same treatment daily and received a total of 11 injections. The route, timing of treatment and the dosages were selected to match prior work, based on ecological relevance, and to span the period of hypothalamic neurogenesis, fetal gonadal development and the early stages of brain sexual differentiation (Arnold & Gorski, 1984; Krishnan, Hasbum, et al., 2019; Krishnan et al., 2018; Krishnan, Rahman, et al., 2019; Rodier, 1980). A subset of offspring (30 male and 29 female) from 9 DMSO, 10 A1221 (PCB), and 10 VIN dams were included in this study. No more than 2 same sex rats per litter were used. We measured body weight and anogenital distance (AGD) on days P7 and P14 to calculate the Anogenital Index 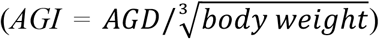. The 5 males and 5 females with the median intrasex AGI measurements were used for the subsequent experiments. The pups were weaned at P21 and re-housed in same-sex groups of 2-3. Beginning on the day of vaginal opening, daily vaginal smears were collected from females and cell cytology was examined as an indication of estrous cyclicity. Timing of pubertal development did not vary across treatment groups (ANOVA; Male age at preputial separation: DMSO 43.50 ± 0.5, PCB 44.22 ± 0.70, VIN 44.10 ± 0.66. Female age at vaginal opening: DMSO 36.44 ± 0.65, PCB 35.30 ± 0.92, VIN 34.60 ± 0.56). Rats were euthanized at ~P120, with females in proestrus, by rapid decapitation and brains removed and processed as described below.

Stimulus Sprague-Dawley rats for the mate preference test were purchased as virgin adults. Males were castrated (GDX) and females ovariectomized (OVX) under isoflurane anesthesia in aseptic conditions (Garcia, Bezner, Depena, Yin, & Gore, 2017; Wu & Gore, 2010). During the surgery, stimulus animals assigned to the hormone-replaced group also had a 1.5 cm Silastic capsule containing testosterone (males: 100% T; GDX+T) or a 1.0cm silastic capsule containing 17β-estradiol (females: 5% E2/95% cholesterol; OVX+E2) implanted subcutaneously into the nape of the neck (Garcia et al., 2017). All rats recovered from surgery for at least one week prior to use in behavioral tests. 32 GDX males (no hormone replacement), 32 GDX+T males, 32 OVX females (no hormone replacement), and 32 OVX+E2 females were used as stimuli throughout the study. For the latter group, on the day of use these E2-treated females were primed for sexual receptivity by a subcutaneous injection of progesterone (P4, 0.6 mg) in sesame oil four hours before experiments started.

### Sociosexual preference behavior

A 1 m x 1 m three-chambered apparatus (Stoelting, Wood Dale, IL) was used as the testing arena (Bell et al., 2016; Reilly et al., 2015). Testing was conducted under dim red light approximately two hours into the dark phase of the light-dark cycle. Each test utilized an experimental (EDC or vehicle exposed) rat at ~3 months of age. Two opposite-sex stimulus rats, one with and one without hormone replacement, were used, with each one placed inside 7 cm x 15 cm cylindrical cages positioned in two far opposite corners of the apparatus. These cages have spaced vertical bars, allowing for limited tactile interactions between rats. The position of stimulus rats was randomized between trials and with respect to hormone status. The bars of the stimulus cage allowed for visual, olfactory, auditory, and minimal tactile interaction between the confined stimulus rat and the freely-moving experimental rat. Each trial began with the two stimulus rats already in position in their cylindrical cages. An experimental rat was placed in the center chamber of the apparatus with closed doors preventing entry into either side chamber for a five-minute habituation period. After habituation, the doors were removed and the experimental rat was allowed to freely explore the entire arena for 10 minutes. Each test was recorded by overhead video. ANY-Maze (Stoelting, Wood Dale, IL) was used to track the position, speed, and distance traveled of the experimental rat in each compartment of the chamber (Hernandez Scudder et al., 2020; Garcia et al., 2017; Reilly et al., 2015). Recordings of the tests were scored by a trained investigator blinded to treatment for the following behaviors: nose touching (direct nose-to-nose contact between the experimental rat and a stimulus rat) and stimulus investigation (all other investigation by the experimental rat of a stimulus rat or stimulus cage). The time the experimental animal spent within one body length of either stimulus cage without engaging with the stimulus animal or cage [time within one body length – (time nose touching + time investigating)] was defined as “time near”. To avoid testing fatigue, stimulus rats were used for no more than three rounds of testing per day and had 10 minutes of rest with access to food and water between each round. Stimulus rats had two days of rest between each day of testing. The entire apparatus was cleaned using 70% ethanol between each test subject.

### Hormone radioimmunoassay

Serum levels of testosterone and corticosterone (CORT) were measured in duplicate samples, and estradiol (E2) in single samples (due to larger serum volume needed for this assay) using radioimmunoassays (Testosterone: MP Biomedicals #07189102, CORT: MP Biomedicals #07120102; E2: Beckman Coulter #DSL-4800). Assay parameters were: CORT, limit of detection 7.7 ng/ml, intra-assay CV 2.5%; testosterone, limit of detection 30 pg/ml, intra-assay CV 3.7%; E2: limit of detection 2.2 pg/ml, intra-assay CV 16.8%.

### TaqMan Low Density qPCR Array

Brains from experimental rats were rapidly removed, chilled on ice, and then coronally sliced at 1 mm using a chilled brain matrix. These slices were placed on slides and stored at -80 until all samples were collected. Bilateral punches were taken of the POA, MeA and VMN using a 1 mm Palkovits punch (Gillette et al., 2014). RNA from frozen POA, MeA, and VMN punches was extracted using AllPrep RNA/DNA Mini Kit (Qiagen, 80204) according to the manufacturer’s protocol. To determine the integrity and purity, a subset of samples was run on a Bioanalyzer 2100 (Agilent, RNA Pico Kit 5067-1513). All samples had a RIN of 8.4 or above. RNA (200 ng) was then converted to single stranded cDNA using high-capacity cDNA reverse transcriptase kit (Life Technologies, 4374966) according to the manufacturer’s protocol. cDNA was run on a custom 48-gene TaqMan Low Density Array Card (ThermoFisher Scientific) with target genes selected based on a priori hypotheses and their role in neuroendocrine function and sensitivity to EDCs reported in the literature. Run parameters were: 95 °C for 10 min, 50 cycles of 95 °C for 15 sec, and 60 °C for 1 min (Topper et al., 2019). Gene expression cycle threshold (Ct) values were normalized using the ΔΔCt method. First, each target gene value was normalized to the expression level of the reference gene *Gapdh* within each subject to generate ΔCt. To standardize between subjects, the Δ Ct of each gene was normalized to the median value of a control group (DMSO females) to generate ΔΔCt. Data are reported as 2^−ΔΔct^. Two genes (*Cyp11a1 & Hsd3b1*) did not amplify and were excluded, leaving 44 target genes and 2 housekeeping genes (*Gapdh, 18s*). In all cases, significance was set at p < 0.05 after appropriate corrections for multiple comparisons.

### Behaviors and hormones

For behaviors, analyses were performed separately for each sex. Those behaviors involving choice based on the hormone status of the stimulus rat were analyzed by a two-way ANOVA (treatment x stimulus hormone status). Other behaviors (e.g. center time, distance traveled of experimental rat) were analyzed by one-way ANOVA. To explore sex differences, a two-way ANOVA for treatment x sex was used for stimulus-independent behaviors between the sexes. Hormone concentrations, body weight, and puberty timing within each sex were analyzed by one-way ANOVA. Reported p-values of multiple comparisons were adjusted using Sidak’s multiple comparisons test.

A hormone preference score was calculated as 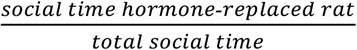. Social preference was calculated as a ratio of time within one body length of both stimulus animals out of the total test time: 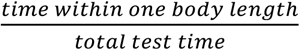. Linear regressions were used to determine correlations with significantly non-zero slopes.

### Principal components analysis

To characterize coordinated phenotypic response to EDC exposure, we performed a Principal Components Analysis (PCA) on morphological, physiological, and behavioral measures including body weight, CORT, E2, T (males only), activity, social preference, hormone preference, and social activity (time spent investigating and interacting with stimulus rats) using the prcomp function in R. Behavioral variables included in the PCA are provided in Table 1. All variables were centered and scaled prior to PCA.

**Table 1.** Behavioral variables analyzed in the sociosexual preference test

|  | Abbreviation<br>in PCA | DMSO |  |  |  | PCB |  |  |  | VIN |  |  |
| --- | --- | --- | --- | --- | --- | --- | --- | --- | --- | --- | --- | --- |
| Behavior |  | Mean | SEM | N |  | Mean | SEM | N |  | Mean | SEM | N |
| **Females** |  |  |  |  |  |  |  |  |  |  |  |  |
| Total distance (m) | Dist | 46 | 3 | 9 |  | 43 | 3 | 10 |  | 43 | 4 | 10 |
| Center time (s) | CentTime | 102 | 14 | 9 |  | 101 | 11 | 10 |  | 118 | 16 | 10 |
| Total social time (s) | SocTime | 335 | 19 | 9 |  | 332 | 22 | 10 |  | 343 | 21 | 10 |
| Total time near stimulus (s) | NearTime | 158 | 11 | 9 |  | 157 | 13 | 10 |  | 163 | 16 | 10 |
| Total nose touch time (s) | NoseTouch | 8.1 | 2.5 | 9 |  | 8.6 | 4.4 | 10 |  | 13.8 | 4.2 | 10 |
| Total stimulus explore time (s) | StimExpl | 169 | 14 | 9 |  | 166 | 28 | 10 |  | 167 | 15 | 10 |
| Hormone preference | HormPref | 0.64 | 0.05 | 9 |  | 0.52 | 0.04 | 10 |  | 0.66 | 0.06 | 10 |
| Social preference | SocPref | 0.56 | 0.03 | 9 |  | 0.56 | 0.04 | 10 |  | 0.57 | 0.03 | 10 |
| **Males** |  |  |  |  |  |  |  |  |  |  |  |  |
| Total distance (m) | Dist | 25 | 2 | 10 |  | 22 | 4 | 9 |  | 23 | 4 | 10 |
| Center time (s) | CentTime | 171 | 34 | 10 |  | 285 | 65 | 9 |  | 317 | 59 | 10 |
| Total social time (s) | SocTime | 256 | 46 | 10 |  | 219 | 52 | 9 |  | **143** | **37** | **10** |
| Total time near stimulus (s) | NearTime | 142 | 37 | 10 |  | 95 | 25 | 9 |  | **68** | **17** | **10** |
| Total nose touch time (s) | NoseTouch | 9.7 | 5.0 | 10 |  | 3.9 | 1.6 | 9 |  | 5.2 | 1.8 | 10 |
| Total stimulus explore time (s) | StimExpl | 105 | 32 | 10 |  | 121 | 38 | 9 |  | 70 | 20 | 10 |
| Hormone preference | HormPref | 0.41 | 0.12 | 10 |  | 0.23 | 0.08 | 9 |  | 0.45 | 0.11 | 10 |
| Social preference | SocPref | 0.43 | 0.08 | 10 |  | 0.37 | 0.09 | 9 |  | 0.24 | 0.06 | 10 |

Sociosexual preference behaviors are shown as mean ± SEM, with n's indicated. Total behaviors are times for the two stimulus rats combined. Bolded numbers are significantly different from the same-sex DMSO control.

### Weighted gene co-expression network analysis

To capture coordinated gene expression changes associated with EDC exposure we performed a Weighted Gene Co-expression Network Analysis (WGCNA) on the 44 target genes measured (Langfelder & Horvath, 2008). WGCNA was performed independently for the two sexes and three brain regions with a minimum module size of five genes. Expression values of each module were summarized as module eigengenes (i.e., the first principal component of each gene co-expression module). Thus, each eigengene is the linear combination of gene expression values that explains the most variation in the expression levels of the genes contained in the module. We assessed coordinated changes in phenotypes and gene expression across treatments using general linear models. Specifically, we tested the hypotheses that the relationship between behavioral measures associated with preference and social interactions and gene co-expression modules describing coordinated nonapeptide gene expression patterns would differ across the treatments in both sexes (Co-expression module ~ Treatment + Behavioral PC + TreatmentXBehavioral PC).

### Co-variance patterns between neural gene expression, physiology, behavior, and morphology

To assess systems-level response to EDC treatment and the potential dis-integration and/or reconstitution, defined as loss or change of correlations by EDCs, respectively, we examined co-variance patterns among all neural gene expression, physiological, behavioral, and morphological measures for each control and treatment conditions separately and visualized changes in the correlation structure across treatments. We calculated Spearman’s rank correlations between all pairwise variables. Variables were clustered using 1-correlation scores as distance variables. To visually assess the extent of integration or re-organization (or lack thereof) for each EDC treatment across levels of biological organization, neural gene expression and phenotype clustering of the control condition was maintained for treatment animals for each sex and brain region.

### Sexually dimorphic neural expression of nonapeptide and sex steroid hormone signaling genes

To characterize changes to sexually dimorphic gene expression and any potential demasculinizing/feminizing or defeminizing/masculinizing effects of EDC treatment, we quantified gene expression distances of all pairwise treatments and sexes for genes from two candidate functional categories, nonapeptides and sex steroid hormone signaling (Fig. 5, yellow and orange functional group, respectively). Nonapeptide genes included *Oxt, Oxtr, Avp, Avpr1a, Kiss1*, and *Kiss1r* for all 3 brain regions, with *Gnrh1* and *Tac3* also included in the POA. Sex steroid hormone signaling genes included *Esr1, Esr2, Ar, Pgr, Nr3c1, Cyp19a1, Srd5a1*, and *Hsd17b1* for all three brain regions. Euclidean distances for each pairwise sex, treatment comparison were calculated using the expression levels of each gene in these functional categories. We then used a permutation analysis to test for significant modifications in sexually dimorphic expression of functional categories. Specifically, to test the hypothesis that sex or treatment were closer in any pairwise comparison than expected by chance, we shuffled sample treatment assignment within sex and recalculated the Euclidean distances for all pairwise comparisons. We repeated this process for 1000 iterations and compared the observed distance to the distribution of permutated distances to obtain a p-value. Distance networks were plotted for both functional categories and all three brain regions.

## Results

### Embryonic exposure to EDCs affected sociosexual behavior in a sex-dependent manner

The mate preference task was performed on 29 females (9 DMSO, 10 PCB, and 10 VIN) and 30 males (10 DMSO, 10 PCB, 10 VIN). Twelve male rats (4 DMSO, 4 PCB, and 4 VIN) failed to investigate both of the stimulus rat options during the allotted 10 minutes. We refer to these males as “non-responders” in all subsequent analyses. All males were included regardless of responder status in analyses of sex differences, PCA analysis, and gene expression analysis. However, Figure 1 shows analyses of only the responder males (6 DMSO, 6 PCB, and 6 VIN), as that test required rats to interact with both opposite-sex stimulus animals to calculate a score. Other figures are inclusive of the entire cohort of males, regardless of responder status.

**Figure 1:**
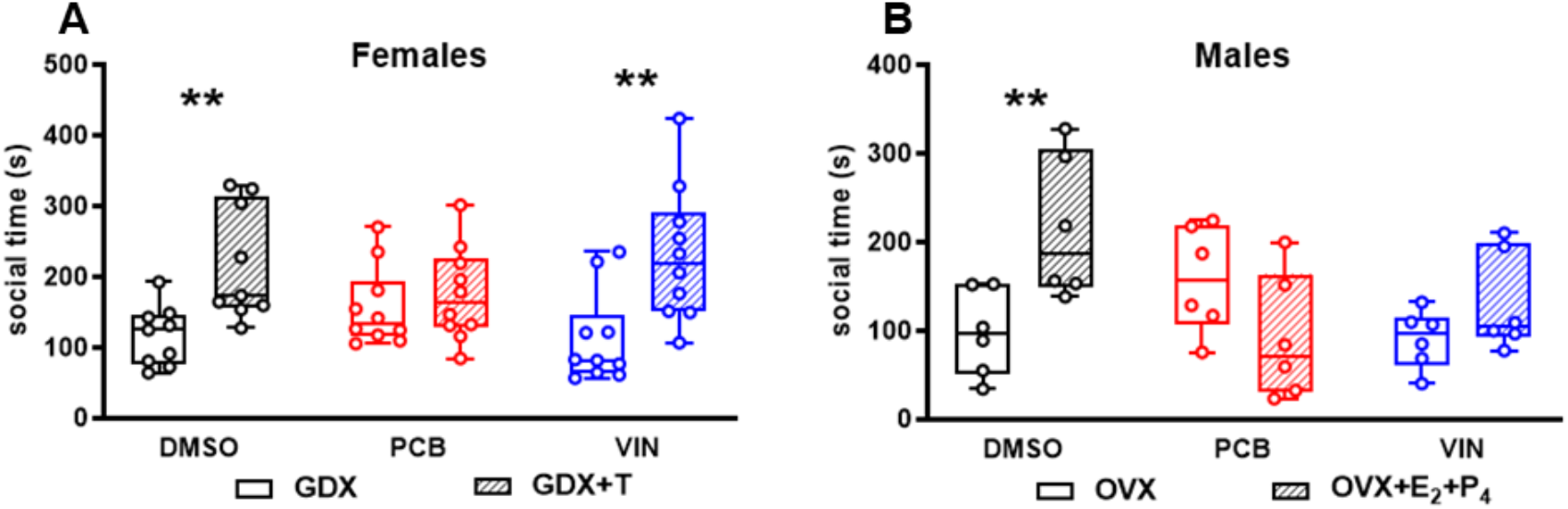
Time spent with each stimulus rat (social time) during mate preference is shown for females (A) and males (B) as median (bar within the box), quartiles (upper and lower limits of the box), and range (whisker) for the 10-minute mate preference test. The same graphing conventions are used for other box-and-whisker graphs. Data for males include responders only. Asterisks indicate a significant difference between the social time spent with the two stimulus options. Two-way ANOVA, main effect of stimulus hormone, followed by Sidak’s multiple comparisons test. GDX, gonadectomized male; OVX, ovariectomized female. ** p <

There was a main effect of hormone status of the stimulus rat on the time experimental females spent associating (within one body length) with the stimulus rats (F_(1, 52)_ = 18.77; p < 0.0001). Females prenatally exposed to DMSO or VIN preferred the hormone-replaced male. Prenatal exposure to PCB, however, abolished this preference in females (Fig 1A). There was no effect of treatment on the total time that the experimental rats spent investigating both stimulus rats (Social Time; Table 1).

In males, there was a significant interaction between treatment and hormone status of the stimulus rat on the time spent associating with the stimulus rats (F_(2, 30)_ = 7.113; p < 0.01). Males exposed prenatally to DMSO spent more time investigating the stimulus female with hormone replacement over the one without. However, the time males exposed prenatally to PCB or VIN spent near the two stimulus rat options did not differ significantly (Fig 1B). These findings replicated those in our recent publication (Hernandez Scudder et al., 2020).

There were significant sex differences in several behavioral measures (Fig 2). Throughout the test duration, females traveled significantly farther than males (Fig 2A; F_(1, 52)_ = 55.94; p < 0.0001). Females spent more time in close proximity to but not interacting with both stimulus rats (Fig 2B; F_(1, 52)_ = 10.49; p < 0.01). Females spent more time directly investigating the stimulus rats than males (Fig 2C; F_(1, 52)_ = 10.43; p < 0.01). For hormone preference score, females preferred the hormone-replaced stimulus animals more strongly than males did (Fig 2D; F_(1, 52)_ = 13.21; p < 0.001). VIN males spent less time in close proximity to the stimulus rats without interacting than DMSO males, shown in Figure 2E as the social preference score (F_(1, 52)_ = 20.66; p < 0.0001). These and all other behavior measures included in further analyses are summarized in Table 1.

**Figure 2:**
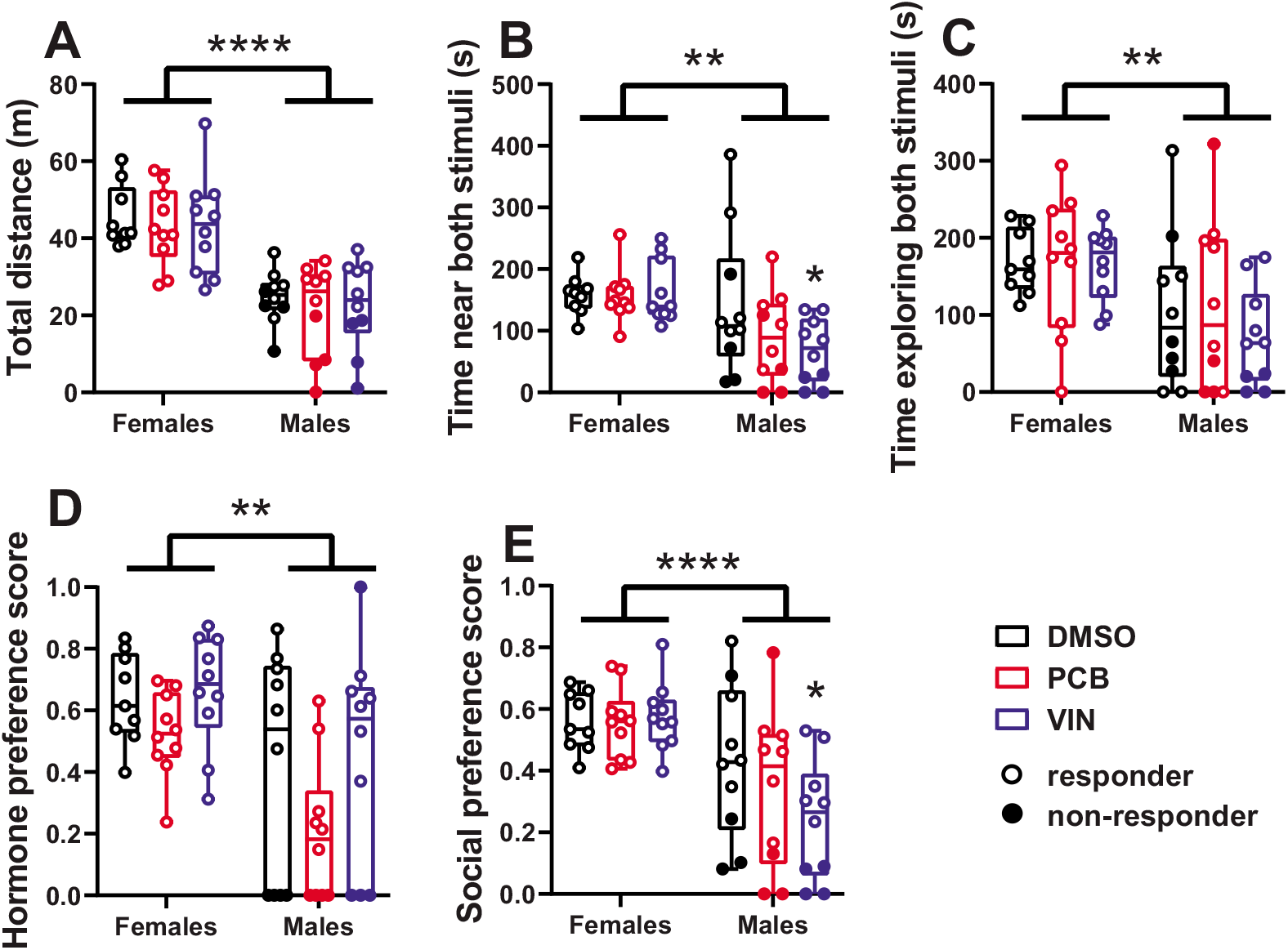
Sex differences during the 10-minute mate preference test are shown. (A) Total distance traveled during the test duration. (B) Total time the experimental rat spent near (within one body length) both stimulus cages without interacting with the stimulus rats. (C) Total time the experimental rat spent exploring (sniffing, touching, etc.) the stimulus cages and stimulus rats (but not nose-touching). (D) Hormone preference score (social time with hormone-replaced stimulus rat/social time with both stimulus rats). (E) Social preference score (social time with both stimulus rats/time in the remote portions of the side chambers [further than one body length away from the stimulus cage]). For males, non-responders (rats who failed to venture near one or both stimulus options) are indicated with solid black circles, here and in subsequent figures. Asterisks indicate a significant sex difference. Twoway ANOVA, main effect of sex followed by Sidak’s multiple comparisons test. ** p < 0.01, **** p < 0.0001, * VIN < DMSO p < 0.05.

### EDCs did not affect circulating steroid hormone levels, but PCBs resulted in reduced body weight in males and females

We measured body weight and serum hormone concentrations (CORT, E2, and T) of behaviorally characterized rats on the day of euthanasia (Fig 3). PCB exposure significantly reduced the body weight of both female (F_(2, 26)_ = 3.506; p < 0.05; Fig 3A) and male (F_(2, 27)_ = 6.080; p < 0.01; Fig 3D) rats. There were no significant effects of treatment on hormone concentrations within males or females [Fig 3B, C (females), E-G (males)]. In females, there was a non-significant trend for PCB exposure to increase E2 levels (F_(2, 26)_ = 2.916; p = 0.07; Fig 3C).

**Figure 3:**
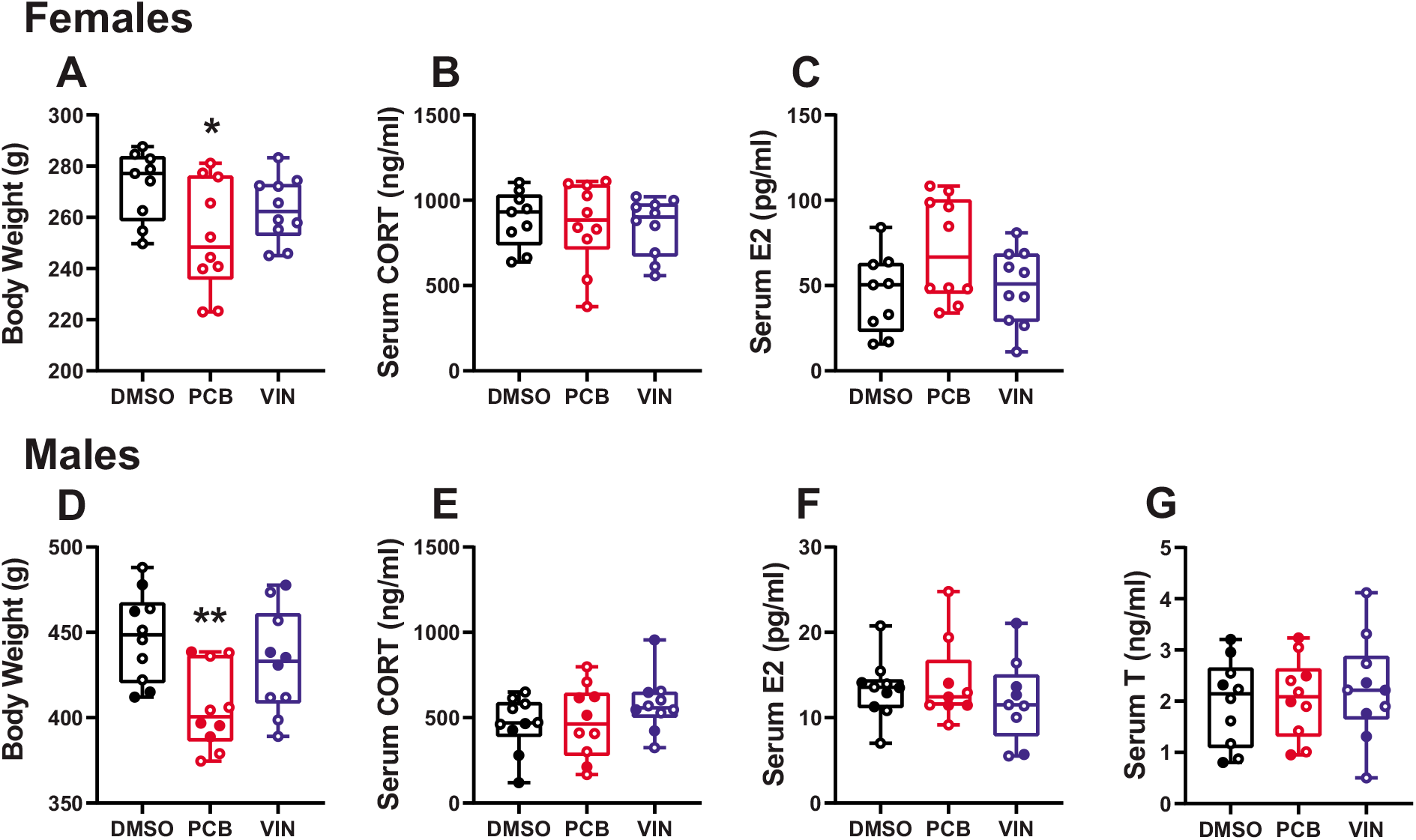
Body weight and serum hormones are shown for female (A-C) and male (D-G) experimental rats. Note differences in y-axes for body weight and serum E2 for the two sexes. Asterisk indicates significant difference from DMSO. One-way ANOVA, main effect of treatment followed by Sidak’s multiple comparisons test. * p < 0.05.

### Integration across levels of biological organization revealed high levels of individual variation with treatment groups

A PCA of behavioral and physiological measures in females (12 measures) and males (13 measures) revealed variation between the sexes in coordinated phenotypic response to EDC treatment, as the PCs characterized different aspects of the phenotype (Fig 4). Eigenvectors describing the loadings, or contribution, of phenotypes to PC variation revealed patterns of coordinated phenotypic variation that differed in strength (absolute value of the eigenvector) and directionality (positive or negative). The results indicate that relationships differed for each treatment group (Fig. 4C and 4D).

In females, the first four PCs described 78% of the total variation in morphology, behavior, and physiology within and across treatments (Fig. 4A). Social Time, Social Preference, and Stimulus Explore loaded strongly and concordantly on PC1 (35%), indicating that PC1 primarily represents variation in time spent engaging in sociosexual behavior. With strong and concordant loadings of Hormone Preference and Near Time, PC2 (20%) represents Social Preference and Social Interaction. PC3 (12%) is strongly loaded by sex steroid (E2) levels and time spent near a stimulus rat with opposing effects. Finally, with strong and opposing loadings of CORT and body weight, PC4 (11% of the variation) may be an indicator of condition. Within treatment, females varied in their integrated response to EDC treatment; however, there were no significant differences across treatments for the first four PCs (Fig. 4E; Supplementary Fig. 1A-F).

In males, the first four PCs described 75% of the total variation in morphology, behavior, and physiology within and across treatments (Fig. 4B). PC1 (38%) primarily described variation among males in the time they spent in the center of the apparatus (Fig. 4D), and, thus, represents differences between responder and non-responder males (Supplementary Fig. 1G. 4F). PC2 (15% of the variation) primarily represented, in opposing fashion, CORT levels and time spent engaging in non-social activity. PC3 (12% of the variation) characterized opposing variation in body weight and T among males. With strong loadings of nose touch, stimulus explore, and hormone preference, PC4 (10% of the variation) characterized variation in preference and social interaction. Because non-responders failed to approach one or both of the stimulus rat options during the allotted 10 minutes biasing their hormone preference and other sociosexual behavioral scores, responder and non-responder males differed across PC4 (Supplementary Fig. 1K).

**Figure 4:**
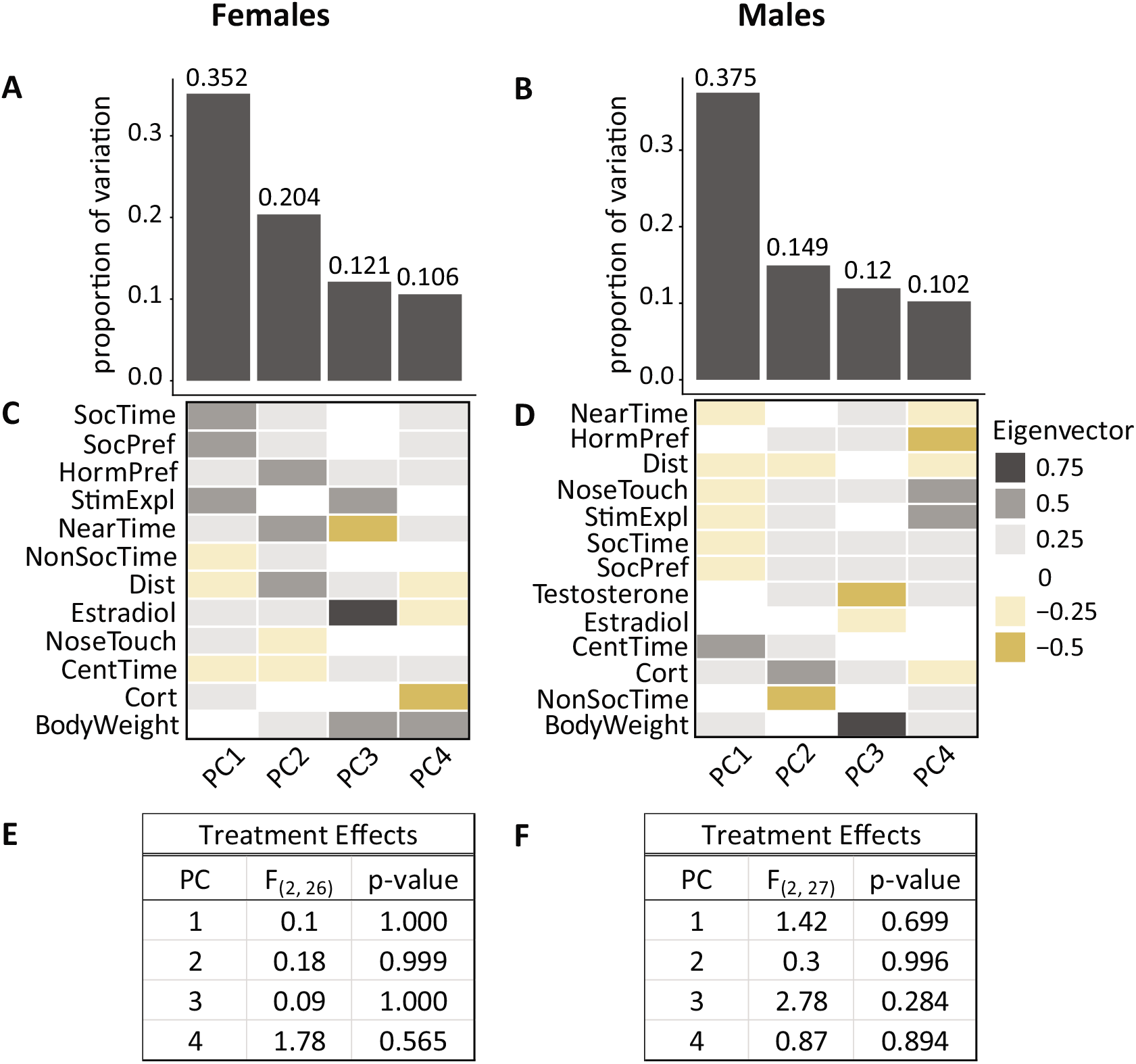
Integrated phenotypic response to EDC treatment. Principal components analysis (PCA) of behavioral and physiological measures in females (left - 12 measures) and males (right - 13 measures) revealed variation between the sexes in coordinated phenotypic response to EDC treatment. The proportion of variation described (A and B) and aspects of the integrated phenotype described by the first four PCs (C and D) are shown. Contribution of each behavioral, morphological, and physiology measures to each PC is indicated by the eigenvectors. Strength of the contribution of each measure is indicated by color saturation. Concordance in directionality of variation among measures is indicated in color. Grays indicate the same direction and browns indicate opposing directions (Fig. C and D). There were no significant treatment differences in either sex for the first four PCs (E and F). Definitions of most included phenotypes are provided in Table 1; in addition, NonSocTime is time spent greater than one body length away from the stimulus rats in the side chambers.

### Embryonic EDC exposure had sex-specific effects on candidate gene expression in VMN, POA, and MeA

The effect of EDC treatment and sex on the expression of the 44 detectable candidate genes in three brain regions was examined (all results shown in Supplementary Table 1). The expression levels of only a small number of genes were significantly affected by treatment after correction for multiple comparisons. In the VMN of females, PCB-exposed rats had higher expression of *Cyp19a1, Oxt, Avp*, and *Kiss1* than DMSO females. In VIN females, *Hsd17b1* and *Oxt* were higher than levels in DMSO females. The expression level of only one gene in the female POA, *Grin2b*, was changed significantly by EDCs: it was over-expressed in the POA of females exposed to PCBs and VIN compared to DMSO. Two genes were affected in the female MeA: *Kiss1* was more highly expressed by PCBs, and *Oxt* expression was lower in PCB treatment and higher in VIN treatment compared to the DMSO control females.

In the male VMN, *Cyp19a1* was lower in both PCB and VIN treatment. In the male POA, only *Grin2b* expression was affected: it was higher in VIN compared to DMSO males. One gene in the male MeA, *Kiss1*, was expressed at lower levels in the VIN treatment group compared to DMSO.

Several sex differences in the gene expression of DMSO rats were identified (Supplementary Table 2). In the VMN, males had significantly higher expression of *Cyp19a1* compared to females. In the POA, females had significantly higher *Kiss1* expression, while males had higher *Grin2b* expression. In the MeA, males had significantly higher *Kiss1* expression than females.

### Gene co-expression modules displayed treatment-specific relationships with preference and social interaction

The 44 candidate genes assessed were selected based on *a priori* hypotheses and their roles in neuroendocrine function and sensitivity to EDCs reported in the literature, as described in greater detail in the Discussion. These genes belong to functional categories that include sex steroid hormone signaling, glucocorticoid stress axis, nonapeptides, GnRH-related genes, neurotrophins, neurotransmission, epigenetics, and clock genes and other transcription factors (Suppl. Table 1; Fig 5). Co-expression modules generated using WGCNA spanned functional gene groups and varied across brain regions and sexes. Treatments did not differ in eigengene expression for either sex or brain region after adjusting for multiple hypothesis testing (Suppl. Fig. 2).

**Figure 5.**
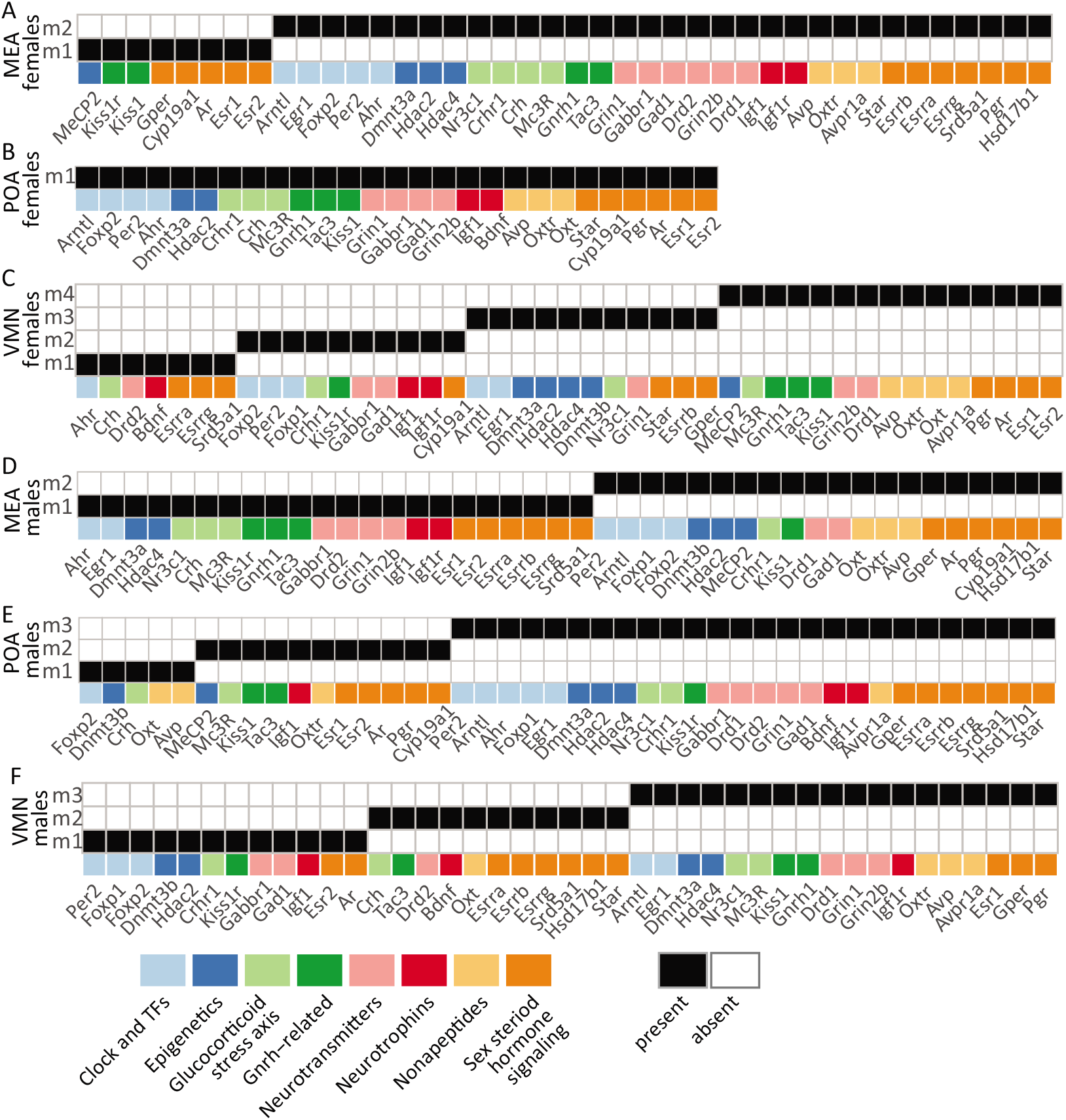
Candidate genes representing eight functional categories were assigned to coexpression modules using WGCNA for each sex (females, A-C; males, D-F) and brain region independently. Genes that did not cluster into any module are not shown. Presence of a gene in a module is indicated in black. Genes are assigned to the functional categories indicated in color. Different number of co-expression modules were identified across the sexes and brain regions. For each sex and brain region, modules are labeled as m1up to m4.

An example of how these data can be used in hypothesis-testing is provided for the nonapeptides. To test the hypothesis that coordinated expression of nonapeptides varies with preference and other social interactions we first identified gene co-expression modules integrating expression of the majority of the nonapeptides including: MeA module 2, POA module 1, and VMN module 4 in females, and MeA module 2 and VMN module 3 in males (Fig. 5). Second, we identified PC2 in females and PC4 in males as representing variation in preference and social interaction across individuals. For PC4 in males, linear models revealed significant treatment and interaction effects in the relationship between module eigengene expression and preference and social interaction in the VMN and a trend in the MeA in males (Fig. 6; Supplementary Table 3).

**Figure 6.**
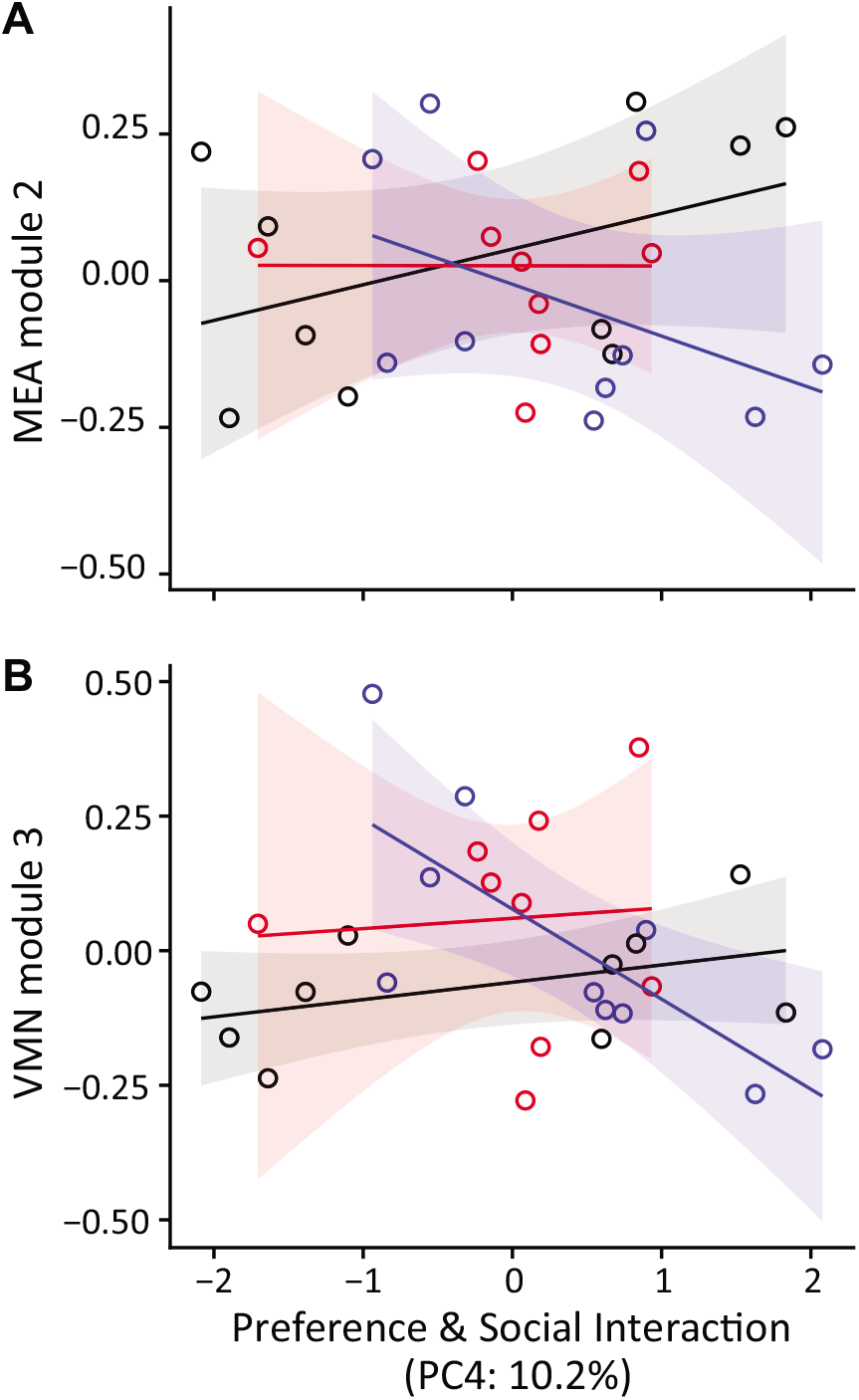
The relationship between expression of gene modules and preference and social interaction differed across treatments in males. Co-expression of gene modules containing nonapeptides genes were compared to principal components describing variation in preference and social interactions in females and males (Supplementary Table 3). In the male MEA a trend highlighting a potential relationship between gene co-expression and preference and social interactions with VIN (blue: t = -0.15; p < 0.1) that is not present in control (black) or PCB (red) treatments (A). In the male VMN there was a significant relationship between gene co-expression and preference and social interactions with VIN (blue: t = -0.20; p < 0.05) that is not present in control (black) or PCB (red) treatments (B). Treatment is indicated in color and shading indicates 95% confidence interval.

### Embryonic exposure to EDCs caused dis-integration and reconstitution across levels of organization

The results presented thus far demonstrate that there was considerable individual variation in both behavior and gene expression. This allowed us to examine co-variance patterns between phenotypic measures (behavior, body weight, hormones) and gene expression to identify any systems-level effects of EDC treatment and ask if behavioral, physiological, and neuromolecular correlations are maintained across treatments. Correlation strengths and phenotypic clustering are illustrated as heatmaps in Figure 7 (in this case for gene expression in the VMN of females). Heatmaps for the POA and MeA in females, and for all three regions in males, are shown in Supplementary Fig. 3). In all cases, we observed that correlations between gene expression levels are stronger (females: median r = 0.3 to 0.43; males: median r = 0.28 to 0.43) than between genes and behavior (females: median r = 0.2 to 0.28; males: median r = 0.2 to 0.32). Importantly, for both sexes and all brain regions the co-variance structure was strongly integrated in control animals (in terms of number and size of robust clusters), whereas with EDC treatment these patterns appeared to “dis-integrate” and/or reorganize. In other words, many genes and phenotypes that were strongly correlated in the control group (indicated as clusters of yellow cells in the heatmap) were uncorrelated or only weakly correlated in treatment groups. To illustrate dis-integration of correlations in the female VMN, the identical order of phenotypes generated by clustering of the DMSO control group (Fig. 7A) was maintained and applied to generate a heatmap for PCB and VIN females (Fig. 7B and 7D), clearly showing few if any robust clusters. If instead the heatmap for PCB and VIN was generated by unbiased clustering, a completely different reconstituted correlation structure emerged (Fig. 7C and 7E), albeit less robust. Importantly, a subset of phenotypes (e.g., *Ahr, Esrrg*, and *Crh* expression are positively correlated in the female VMN across treatments, Fig. 7) maintained strong correlations across all treatment groups indicating their robust relationships. While the present study is underpowered to perform a quantitative analysis of this “dis-integration hypothesis,” the pattern of correlation loss and reorganization in both sexes and in all three brain regions reveals striking qualitative differences between the DMSO and EDC-treated rats.

**Figure 7:**
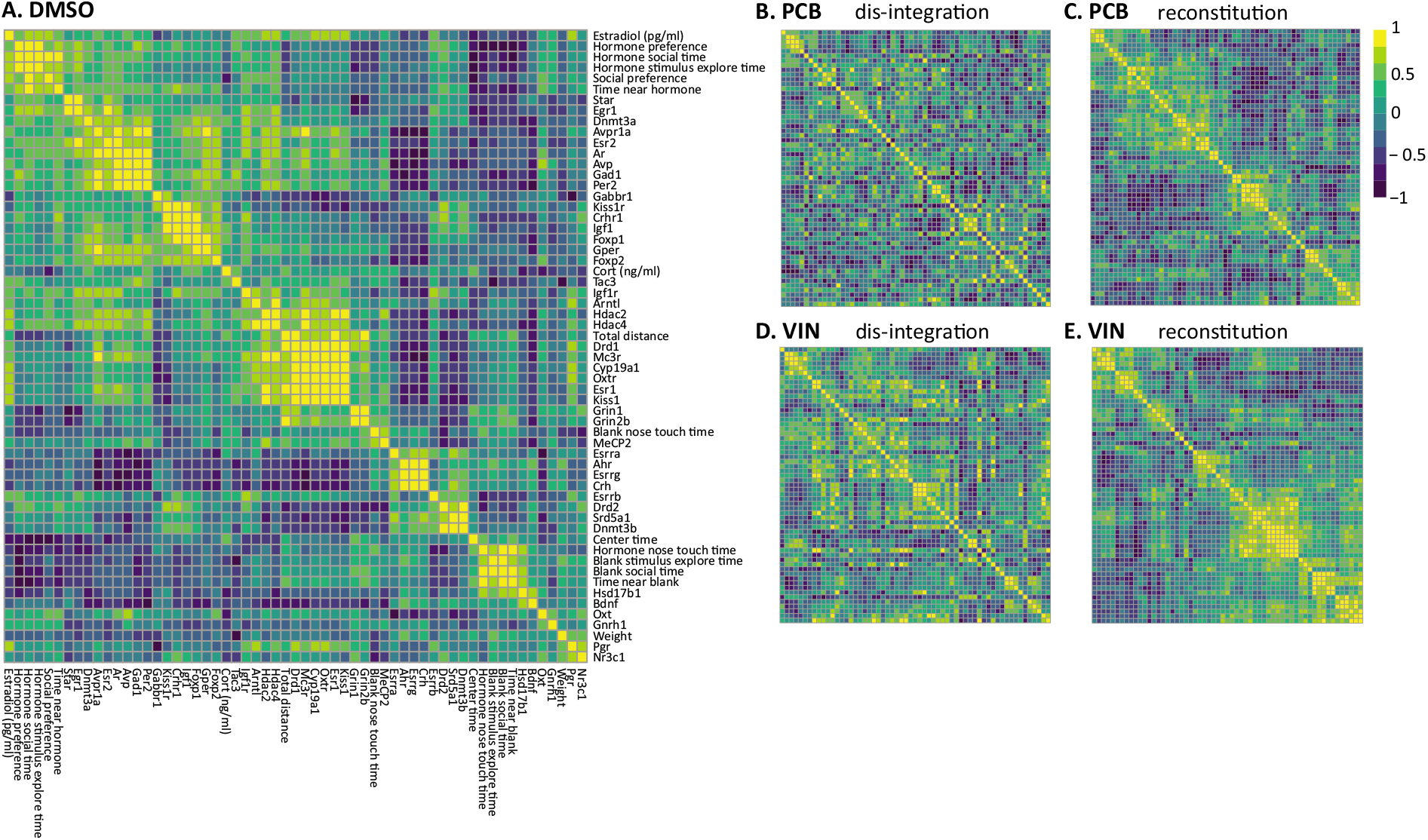
Representative correlation heatmap from the VMN of females, illustrating the dis-integration and reconsolidation of behavioral, hormonal, and neuromolecular phenotypes caused by EDC treatment. Gene expression, behavioral, and physiological measures were clustered using Spearman’s rank correlations for the DMSO control samples (A). The same ordering of phenotypes in the control group was applied to both the PCB (B) and VIN (D) treatment groups to illustrate dis-integration associated with EDC treatment. To illustrate reorganization of phenotypes, behavioral, hormonal, and neuromolecular phenotypes were clustered for using Spearman’s rank correlations for PCB (C) and VIN (E) females. Correlation strength is indicated by intensity of color with yellow indicating positive correlations and indigo indicating negative correlations. Correlations of remaining brain regions and sexes including the order of phenotypes in the reorganized heatmaps above are provided in Supplementary Fig. 3.

One set of factors that demonstrates this dis-integration effect particularly well is the total distance traveled during the mate preference test and VMN Grin2b expression in females (Fig 8). A robust correlation between these factors was found for the DMSO group (F_(1,7)_ = 19.60; p < 0.01; R^2^ = 0.74; Fig 8A) but not the PCB (R^2^ = 0.005; Fig 8B) or VIN (R^2^ = 0.0008; Fig 8C) females. Conversely, we found strong evidence of reconstitution in males, where the direction of correlations present in the DMSO group were reversed by EDC treatment. Fig 8D shows one such example: the negative correlation between hormone preference score and *Hsd17b1* expression in the POA of DMSO males (F_(1,8)_ = 13.56; p < 0.01; R^2^ = 0.63). The direction of this correlation was reversed by PCB treatment (F_(1,7)_ = 17.68; p < 0.01; R^2^ = 0.72; Fig 8E) and abolished by VIN treatment (R^2^ = 0.03; Fig 8F). Furthermore, there was considerable individual variation in both gene expression and behavior, and the correlation heatmaps demonstrate that there are many stronger correlations between genes than between genes and behavior.

**Figure 8:**
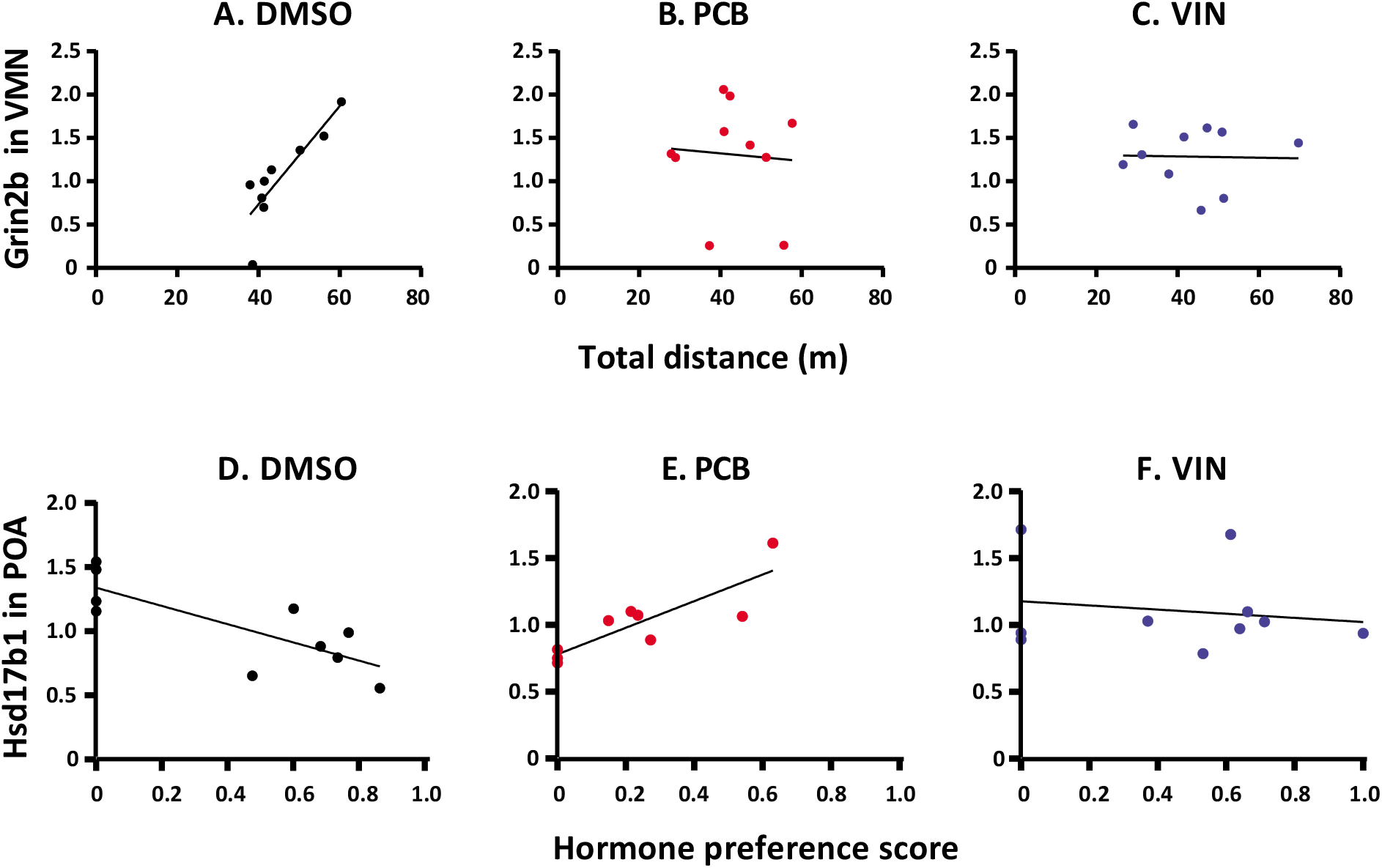
Example of treatment-induced dis-integration and reconstitution of behavioral measures and gene expression. Correlation between (A) total distance traveled during the mate preference test and Grin2b expression in the VMN of female DMSO rats. (B) No correlation between measures after PCB exposure. (C) No correlation between measures after VIN exposure. (D) Negative correlation between hormone preference score and Hsd17b1 expression in the POA of male DMSO rats. (E) Positive correlation between hormone preference score and Hsd17b1 expression in the POA of male PCB rats. (F) No correlation between measures after VIN exposure.

### EDC exposure disrupts sexually dimorphic expression of nonapeptides and sex steroid hormone signaling genes

To test the hypothesis that EDC treatment interferes with sexually dimorphic gene expression patterns, or, based on the behavioral phenotype, specifically, demasculinizes (VIN, PCB in males) or defeminizes (PCB in females) neural gene expression patterns of our experimental rats, for each brain region, we calculated the Euclidean distance in expression for all pairwise sex, treatment comparisons for genes from two functional categories, nonapeptide and sex steroid hormone signaling genes (from Fig. 5, yellow and orange groups, respectively). While no generalized demasculinizing or defeminizing effects of EDC treatment was evident across all brain regions (Suppl. Figure 4), several interesting patterns emerged. In the MeA, modification of sexually dimorphic nonapeptide and sex steroid hormone signaling gene expression in response to EDC treatment was quite similar (Fig. 9). For both nonapeptides and sex steroid hormone signaling genes, males treated with VIN and females treated with PCB were significantly more similar to each other than expected by chance (Fig. 9 – line between red circle and blue triangle). This finding indicates that EDC treatment shifted both sexes away from their respective control groups and toward an intermediate defeminized (females) and demasculinized (males) phenotype. In addition, males treated with PCB were significantly more similar to control males than expected, indicating a muted response of nonapeptide and sex steroid signaling in the MeA to this treatment (Fig. 9).

**Figure 9.**
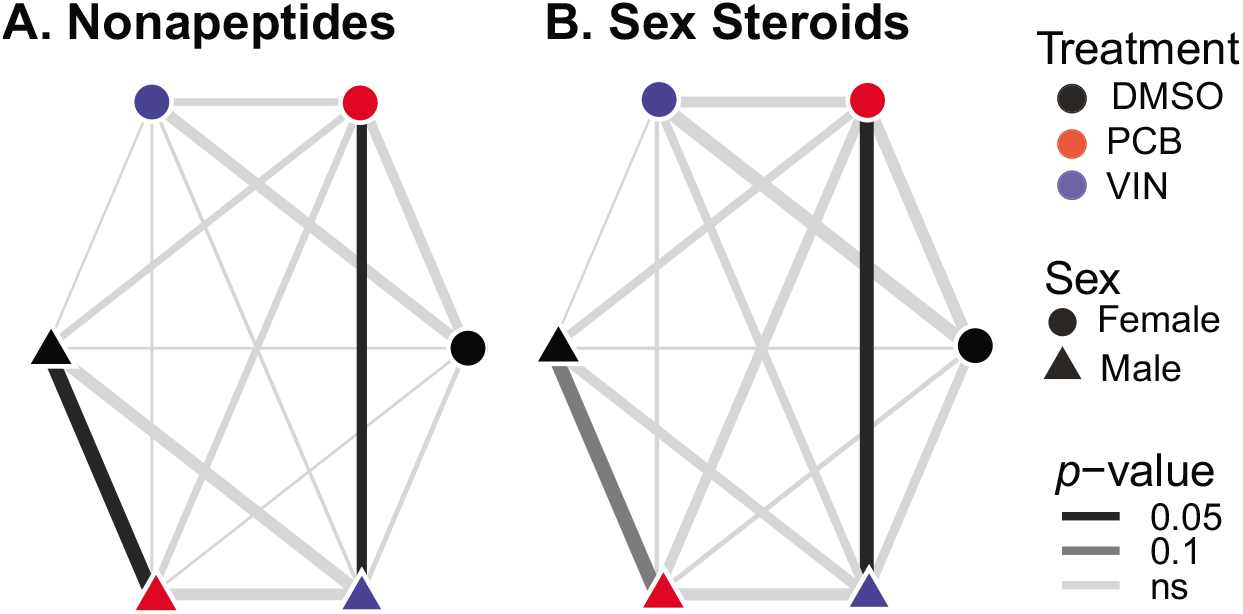
MeA distance networks indicate gene expression distance for all pairwise sex and treatment comparisons for nonapeptide (A) and sex steroid hormone signaling genes (B). Nodes (circles, female; triangles, male) represent each sex and treatment. Edge width is defined as – z-score of all pairwise Euclidean distance in the network such that thick edges represent nodes that are more similar in gene expression. Darker edge colors indicate that nodes are closer than would be expected by chance. Distance networks of remaining brain regions are provided in Supplementary Fig. 4.

## Discussion

The current study a novel analysis of the changing phenotypic relationships among neural gene expression and behavior, and the dis-integration and reconstitution among related sociosexual behavioral measures and neuromolecular networks, in rats exposed prenatally to EDCs. More specifically, sociosexual preference behaviors were impaired by PCBs in both sexes, and by VIN in males. This difference between the EDCs is interesting and may reflect the different modes of action by which the PCB mixture, A1221, acts [mainly weakly estrogenic (Dickerson & Gore, 2007)] vs. that of VIN [anti-androgenic (Euling et al., 2002; Stroheker et al., 2005)]. These results are also consistent with the literature showing that phenotypes induced by EDCs are compound-specific, likely reflecting different mechanisms by which a compound can disrupt the endocrine system. In line with the current finding, Crews et al. (Crews et al., 2012) demonstrated the utility of using an approach to combines levels of biological organization to produce the ‘functioning phenotype’ in a model of transgenerational exposures using VIN. Here, each of the EDCs (PCBs, VIN) resulted in a unique dis-integration/reconstitution of the behavioral and molecular relationships within each sex. As a whole, the perturbations by EDCs of conspecific interactions have implications for social preference and sexual selection (Gore, Holley, & Crews, 2018).

### Prenatal EDCs changed mate preference in a treatment- and sex-specific manner

Female mammals have an incentive to seek out mates with the best likelihood of producing fertile offspring due to this sex’s high investment in reproduction. For example, males with the typical adult range of concentrations of testosterone (Spiteri et al., 2010), and odors from a male with higher testosterone levels, are preferred by sexually active female rats over their low- or no-testosterone counterparts (Osada, Kashiwayanagi, & Izumi, 2009; Taylor et al., 1982). Here we showed that prenatal exposure to PCBs abolished the females’ preference for a stimulus male with testosterone replacement over a male without testosterone, replicating results from a recently published study (Hernandez Scudder et al., 2020). This outcome could translate into compromised reproductive success if a female were unable to discriminate between optimal and sub-optimal males in more naturalistic conditions. Interestingly, VIN treatment had no effect in the females. The difference between the EDCs may be attributable to differential hormonal mechanisms acted upon by the different classes of EDCs (Nugent et al., 2015; Schwarz & McCarthy, 2008).

While males tend to be less choosy about mates, the process of mating involves mutual interactions and coordination between both members of the dyad. Females exhibit proceptive behaviors to solicit the sexual attention of males, and males are also able to discriminate the odor of urine from receptive females (Edwards & Einhorn, 1986; Hurtazo, Paredes, & Ågmo, 2008; Lydell & Doty, 1972; Xiao, Kondo, & Sakuma, 2004). In the current study, and unlike females, exposure of experimental males to both classes of EDCs (PCB or VIN) abolished the preference for the hormone-primed female over the female without hormone-replacement. The male rat brain develops under the influence of relatively high concentrations of both androgens and estrogens (Bakker et al., 2006; Nugent et al., 2015; Schwarz & McCarthy, 2008; Wright et al., 2010), perhaps conferring greater sensitivity to disruption of these pathways by both VIN and PCBs, respectively.

Previous work on prenatal A1221 exposure showed disrupted sex behavior in female rats, and decreased sexual motivation in male rats (Steinberg et al., 2007; Topper et al., 2019). Exposure to other PCBs caused reduced sexual motivation and receptivity in females and altered sexual behavior in males (Colciago et al., 2009; Cummings et al., 2008; Faass et al., 2013; Faqi, Dalsenter, Merker, & Chahoud, 2016; X. Q. Wang, Fang, Nunez, & Clemens, 2002). Exposure to VIN during the prenatal period and during postnatal life (E14 to adulthood) resulted in a lack of sexual motivation and deficits in sexual performance (reduced erections and ejaculations) in male rabbits (Veeramachaneni, Palmer, Amann, & Pau, 2007; Veeramachaneni et al., 2006). Our current finding adds to this literature on sex-specific effects of EDCs on sociosexual behaviors. Subsequent integrative and systems-level analyses provide novel insights into patterns of dis-organization and reconstitution into novel phenotypes.

### Prenatal EDCs altered the neuromolecular phenotype in the hypothalamus and amygdala

Prenatal EDCs affected the expression of a small number of genes in the VMN, POA, and MeA. It is notable that genes for kisspeptin (*Kiss1*), nonapeptides (*Avp, Oxt*), steroidogenic enzymes (*Cyp19a1, Hsd17b1*), and the glutamatergic NMDA receptor subunit 2b (*Grin2b*) were those affected, as these same genes have previously been shown to be disrupted by EDCs and are involved in sexually-dimorphic behaviors. For example, the hypothalamic kisspeptin system is highly estrogen sensitive (Jenny Clarkson, Boon, Simpson, & Herbison, 2009; Navarro & Tena-Sempere, 2008), making it an obvious target for estrogenic EDCs. Other studies have shown that PCBs and other estrogenic EDCs affect kisspeptin protein and gene expression (Cao, Mickens, McCaffrey, Leyrer, & Patisaul, 2012; Dickerson, Cunningham, Patisaul, Woller, & Gore, 2011; Ruiz-Pino et al., 2019), consistent with the current results. Our finding that VIN affects *Kiss1* in males is also consistent with this neuropeptide’s regulation by androgens (Cernea, Phillips, Padmanabhan, Coolen, & Lehman, 2016; Clarkson, Shamas, Mallinson, & Herbison, 2012).

Oxytocin gene expression was decreased in the MeA of PCB females; this was the same group of rats that showed deficits in the mate preference test. The MeA has been characterized as an important target of oxytocin in mate and odor preference behaviors (Yao, Bergan, Lanjuin, & Dulac, 2017), but, although it has relatively sparse oxytocin fibers, there is evidence for a role of oxytocin expression in the amygdala in social behavior (Smith, DiBenedictis, & Veenema, 2019). Oxytocin knockout mice have deficits in social recognition associated with reduced activity in the MeA and its projection targets (Ferguson, Aldag, Insel, & Young, 2001). By contrast, *Oxt* was increased by both PCBs and VIN in the female VMN in the current study, and vasopressin by PCBs in the female VMN. Other labs have reported effects of EDCs on the nonapeptides vasopressin, oxytocin, and their receptors in several brain regions (Arambula, Jima, & Patisaul, 2018; Sullivan et al., 2014; Witchey, Fuchs, & Patisaul, 2019), implicating these as targets for perinatal endocrine disruption.

Both PCB and VIN males had lower *Cyp19a1* (aromatase) expression in the VMN compared to DMSO control males. This region exhibits some of the highest levels of aromatase in the brains of rats along with the POA and the bed nucleus of the stria terminalis (BNST) (Wagner & Morrell, 1996). In the hypothalamus, aromatase expression and activity is sexually dimorphic with males having denser expression and higher activity (Roselli, Klosterman, & Fasasi, 1996). Prenatal exposure to a similar PCB, Aroclor 1254, reduced aromatase activity in the hypothalamus of neonatal male rats (Hany et al., 1999). Prenatal exposure to another EDC, the phthalate DEHP, reduced *Cyp19a1* expression in the hypothalamus of neonatal rats (Gao et al., 2018).

The N-methyl-D-aspartate (NMDA) glutamate receptor subunit 2b (*Grin2b*) is expressed widely throughout the rat hypothalamus (Eyigor, Centers, & Jennes, 2001) and its presence and abundance affects functional properties of NMDA receptors. In the POA, PCBs resulted in the over-expression of *Grin2b* in both sexes, and VIN also increased *Grin2b* in the female POA. Hypothalamic *Grin2b* expression is sensitive to circulating estradiol levels, and naturally decreases during reproductive senescence (Maffucci, Noel, Gillette, Wu, & Gore, 2009). The activation of gonadotropin-releasing hormone (GnRH) neurons in the POA by glutamate is necessary for reproductive function, and administration of a specific antagonist of the NMDAR2b subunit altered GnRH and downstream luteinizing hormone (LH) release in rats (Maffucci, Walker, Ikegami, Woller, & Gore, 2008). Limited work also suggests that EDCs may change *Grin2b* expression (Alavian-Ghavanini et al., 2018; Dickerson, Cunningham, Patisaul, et al., 2011). Our finding suggests that glutamatergic neurotransmission may be altered by prenatal EDC exposure.

While we found no evidence for globally demasculinizing or defeminizing effects of EDC treatment on gene expression, the typically sexually dimorphic expression of nonapeptide and sex steroid signaling genes was significantly disrupted in the MeA. Specifically, we found that EDC treatment (VIN males and PCB females) shifted both sexes away from their respective control groups and toward an intermediate defeminized (females) and demasculinized (males) phenotype (Fig. 9). Further work using global gene expression profiling, something we are in the midst of undertaking, will enable us to better test this hypothesis. Some evidence is provided by a previous study from our lab (Walker, Goetz, & Gore, 2014) in which a shorter-term (2 day) prenatal exposure to PCBs changed gene expression patterns in the anteroventral periventricular nucleus of the hypothalamus (AVPV) in female rats such that developmental profiles were masculinized. Furthermore, hierarchical clustering analysis of genes in the AVPV and the arcuate nucleus revealed changes in relationships among gene expression profiles, with males and females each being affected in a sex-specific manner. That prior study’s results presaged those of the comprehensive analyses of the current one and its finding of dis-integration and reconstitution, as discussed next.

### EDC treatment can dis-integrate and/or reconstitute the relationships of behavior and gene expression phenotypes

Gene co-expression patterns revealed sex, brain region, and treatment-specific effects of EDCs. Genes clustered into distinct modules between males and females across all brain regions suggesting sex-specificity of neuromolecular phenotypes. In VIN males there was a relationship between gene co-expression of modules containing nonapeptides and the behavioral PC describing variation in social interaction and preference in the VMN (and a trend in the MeA) that was not present in DMSO or PCB males. This finding indicates that a reorganization of the neuromolecular and behavioral phenotypes occurs in males in response to prenatal VIN exposure. Previous studies have reported effects of EDCs on the nonapeptides vasopressin, oxytocin, and their receptors in several brain regions (Arambula et al., 2018; Sullivan et al., 2014; Witchey et al., 2019), implicating these as targets for perinatal endocrine disruption. Interestingly, gene co-expression modules clustered genes across functional categories and often clustered genes from the same functional category into distinct modules (see Fig. 5). This finding illustrates the importance of a systems-level approach that describes the neuromolecular phenotype beyond typical candidate pathways to identify gene expression mechanisms of complex behavior.

Furthermore, there were many more correlations among the expression of genes rather than between genes and behavior. The correlation heatmaps showed a striking pattern of EDC influence on these relationships. In both males and females, we observed an EDC-induced qualitative dis-integration of gene-gene and gene-behavior interactions in all three brain regions.

## Conclusions

Overall, this present study presents evidence that prenatal exposure to two classes of EDCs abolished the innate preference of males and females for an opposite-sex mate, and identified several gene targets modified by EDC treatment both independent of and related to specific behavioral measures. More importantly, the relationships across the different levels of phenotypic analysis underwent considerable dis-organization and reorganization, indicating that beyond effects on individual genes and behaviors, EDCs disrupt the integration across these levels or organization. Because there were stronger correlations between genes than there were between genes and behavior, a broader analysis of more genes and brain regions is merited. These findings provide a foundation for further work on the disruption of complex behaviors after prenatal exposure to EDCs.

## Acknowledgments

The authors thank Dr. Krittika Krishnan and Dr. Michael Reilly for their assistance collecting behavioral data and for their input on the selection of target genes.

---

**Supplementary Figure 1.**
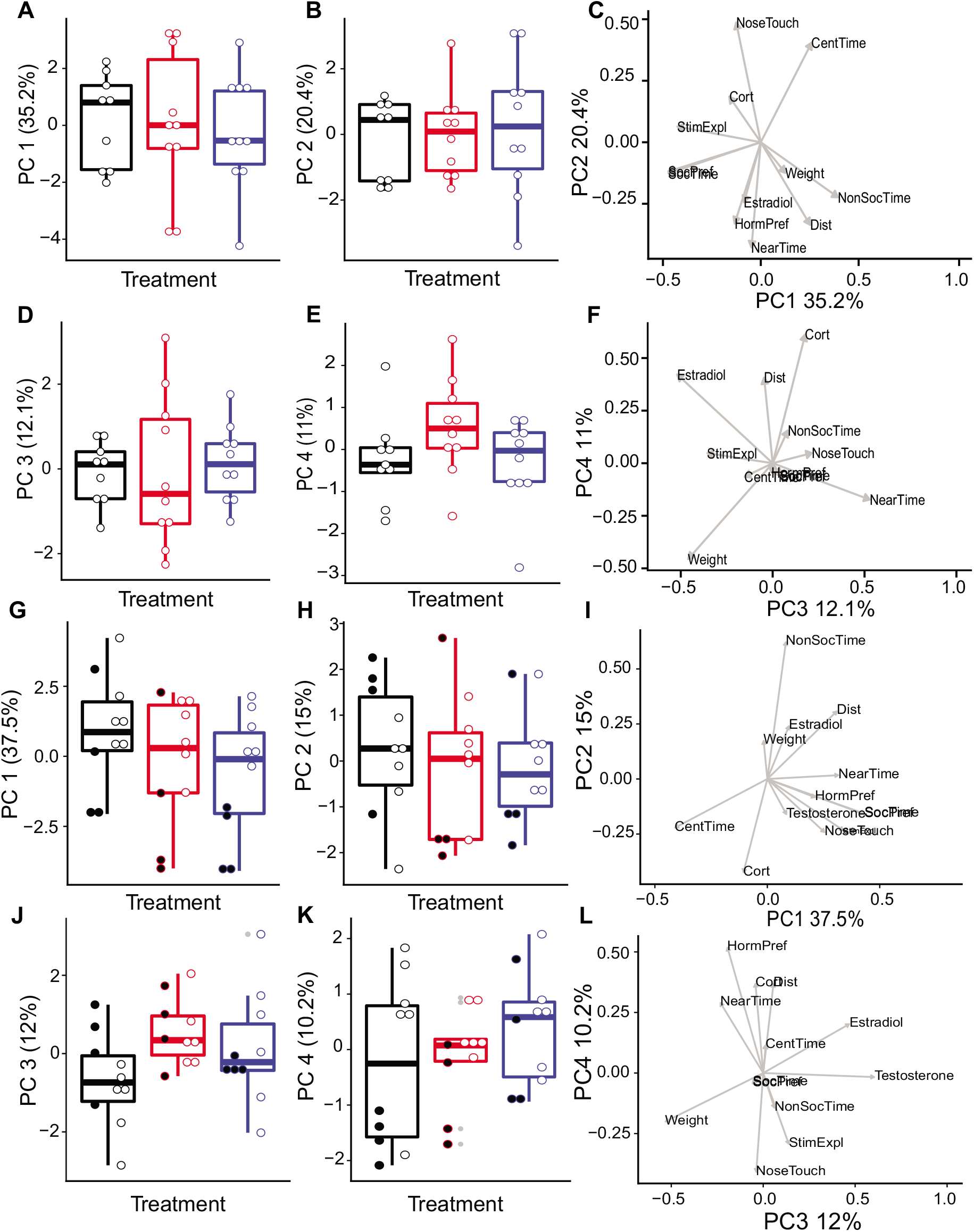

**Supplementary Figure 2.**
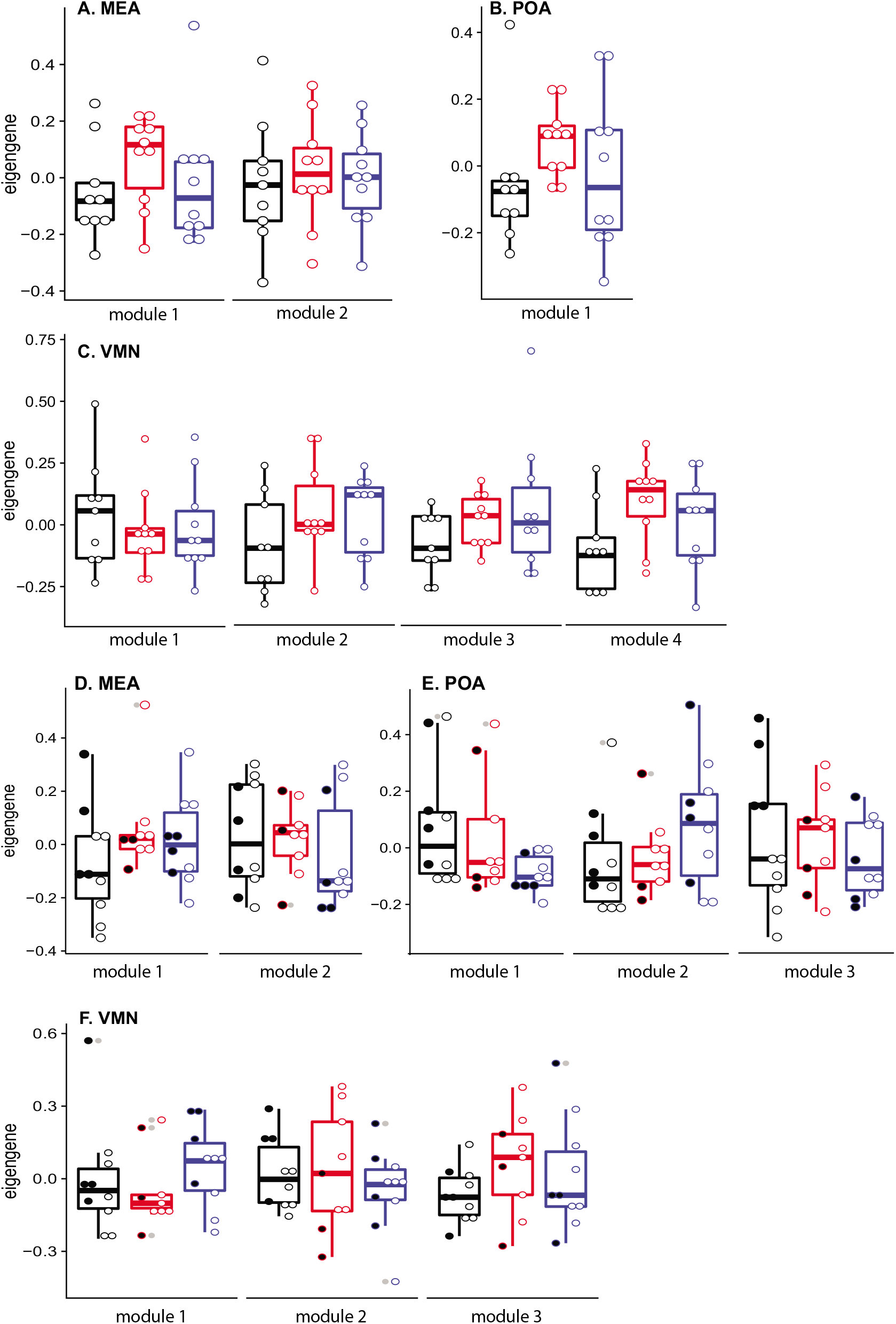

**Supplementary Figure 3.**
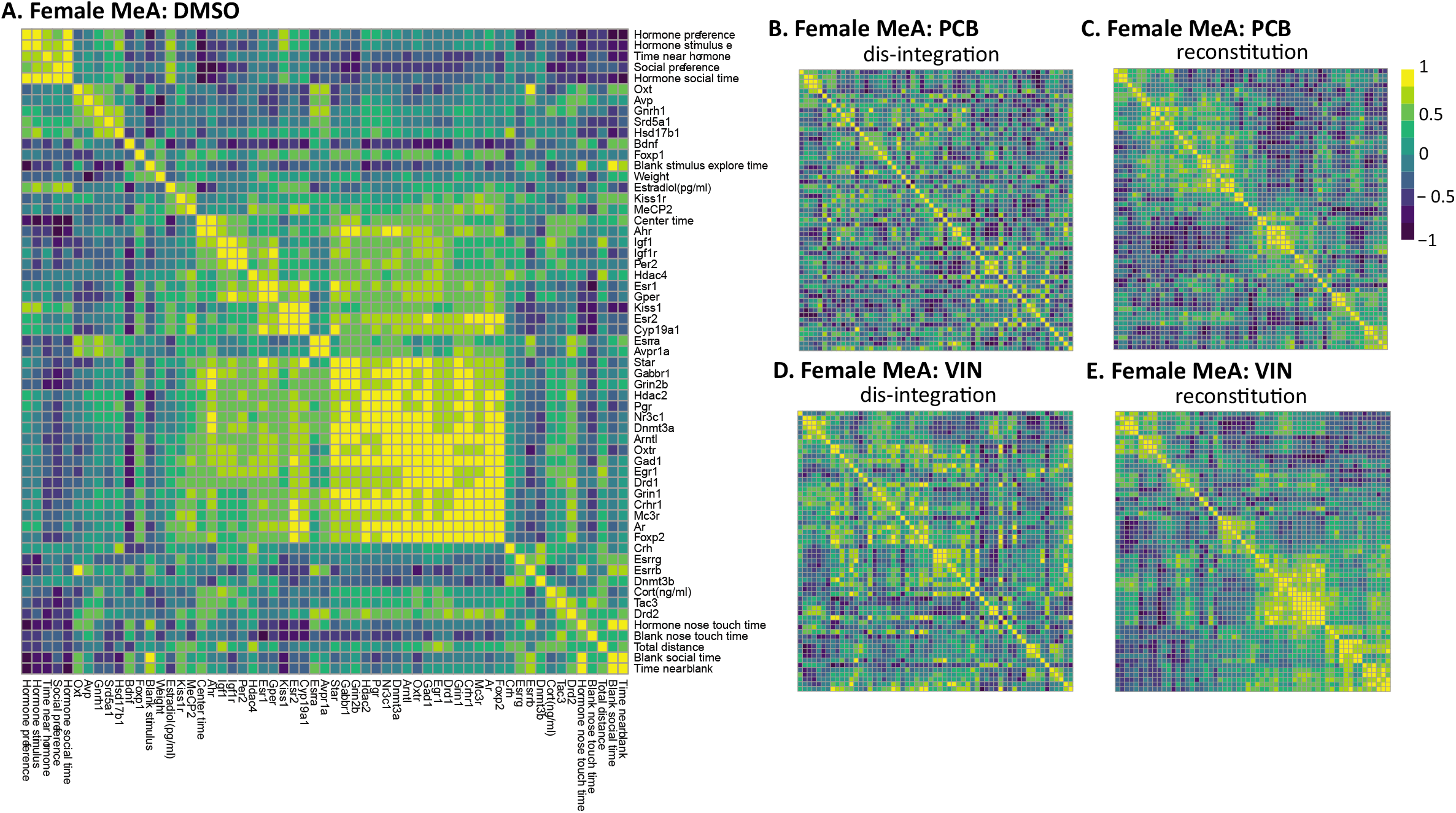
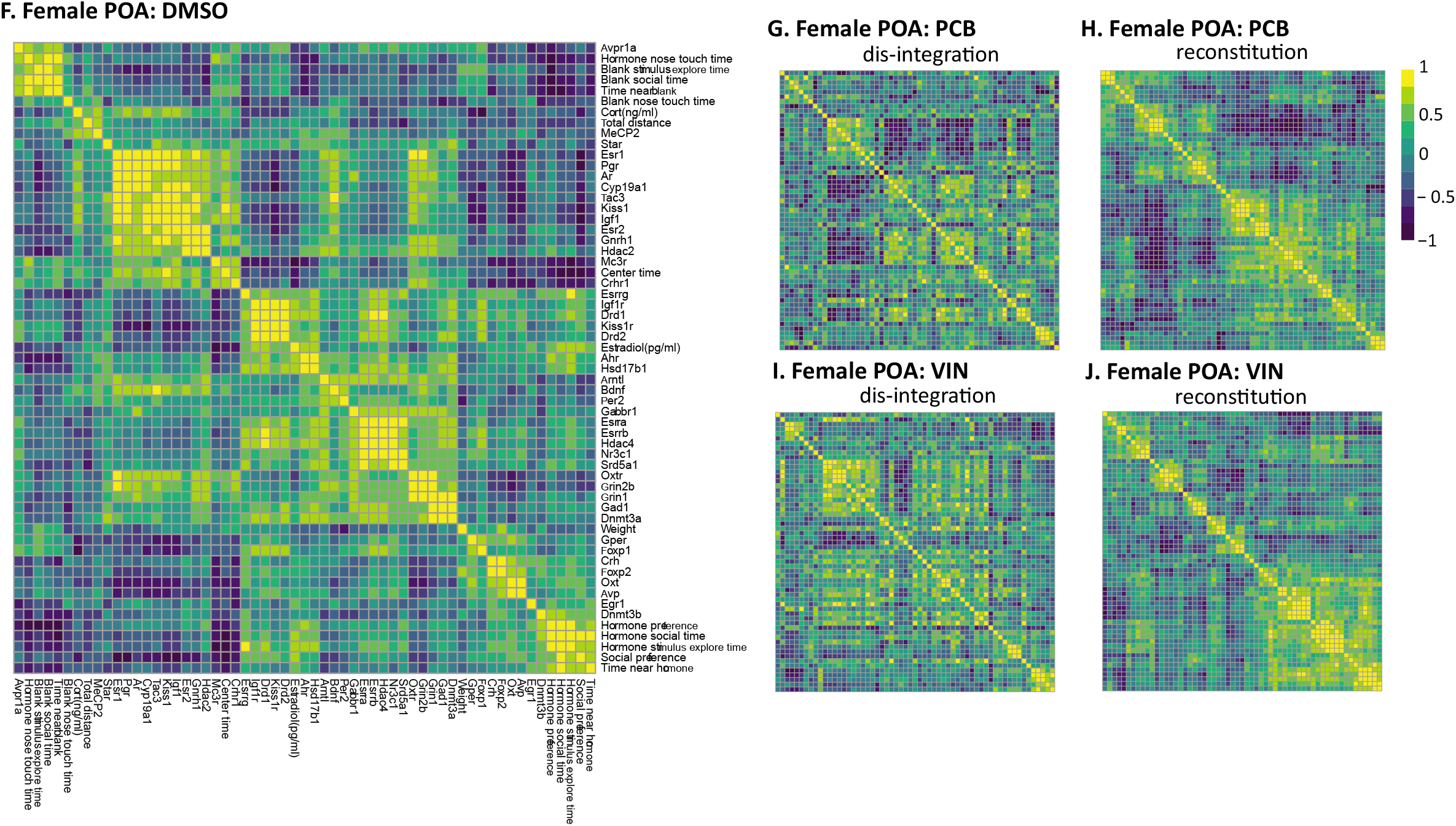
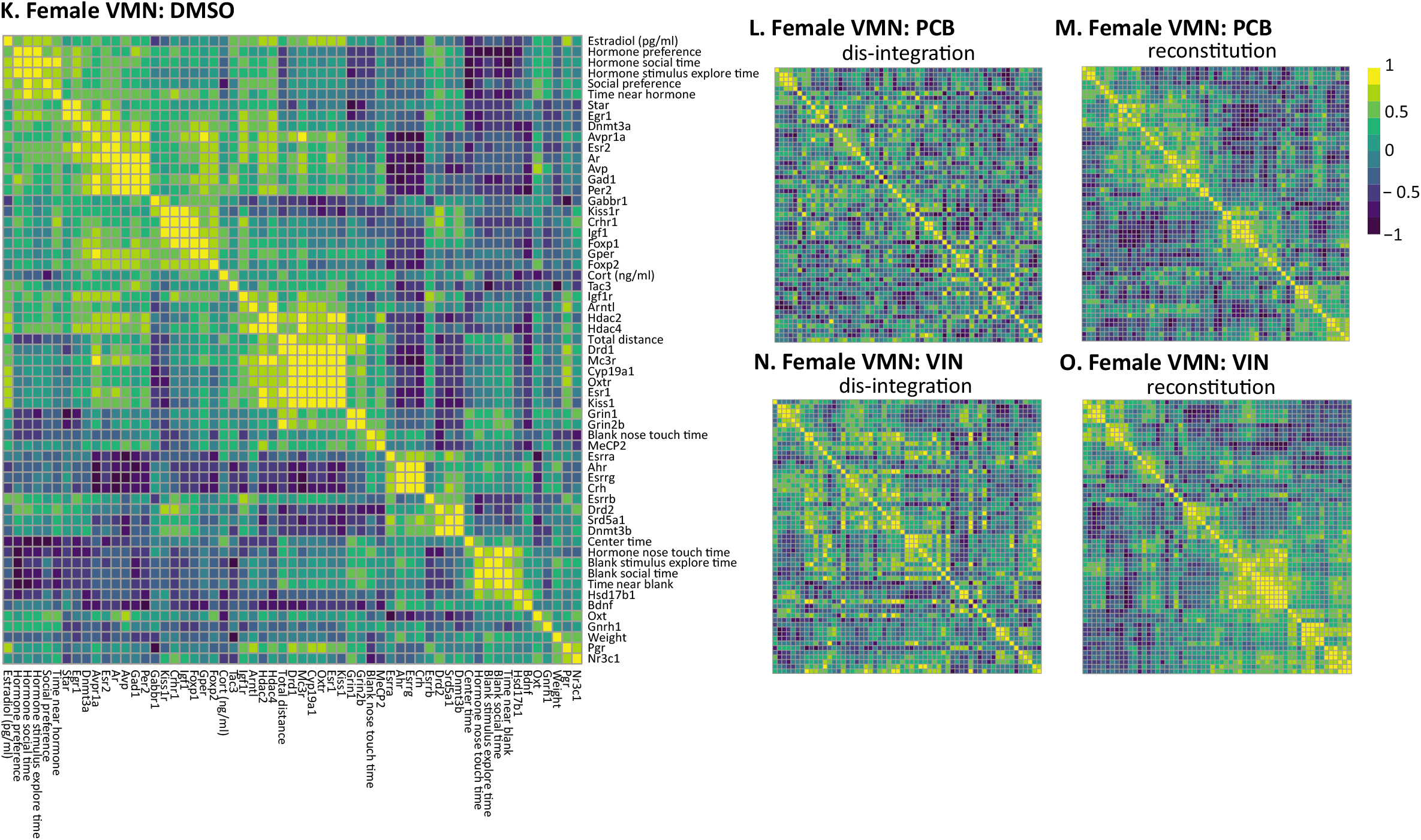
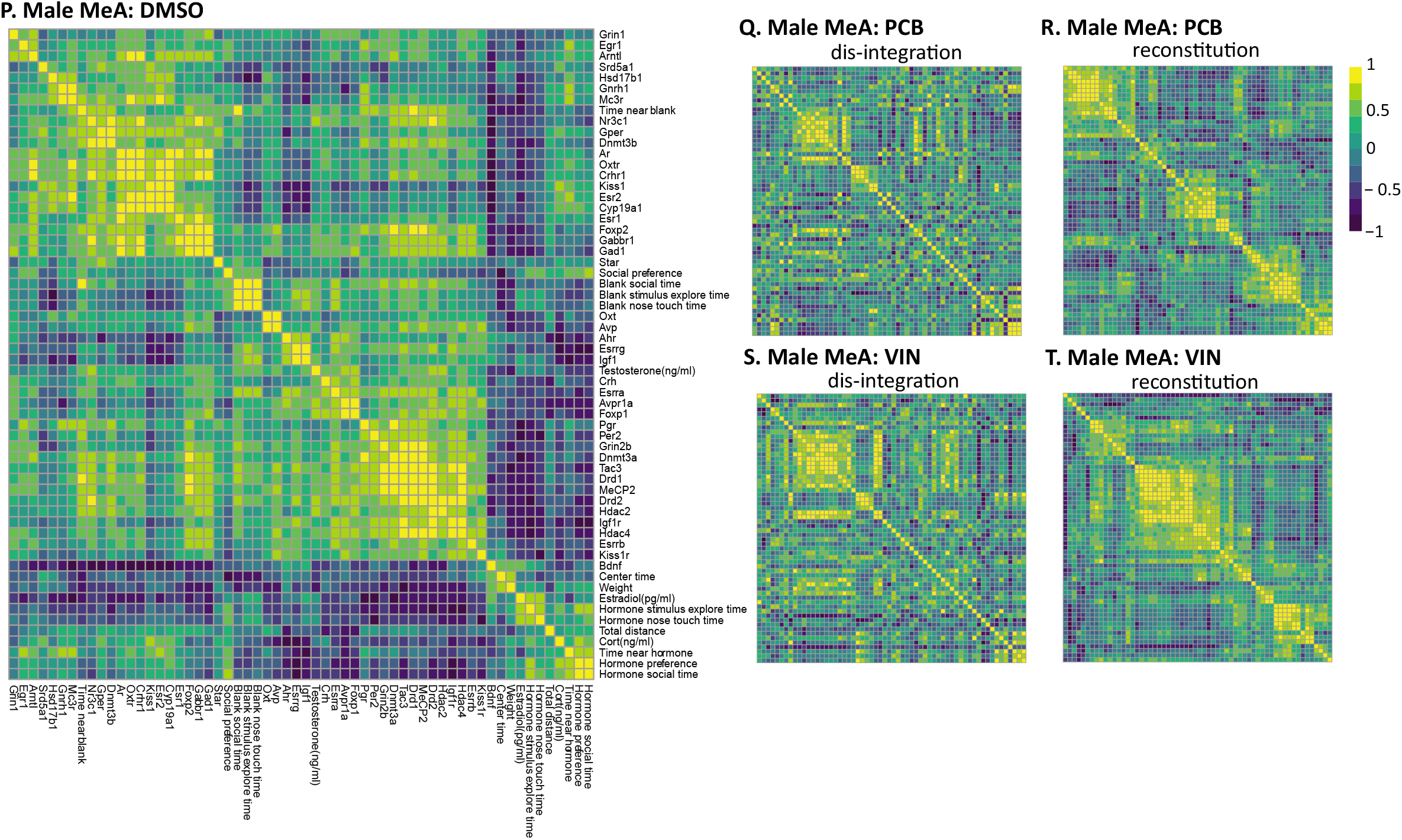
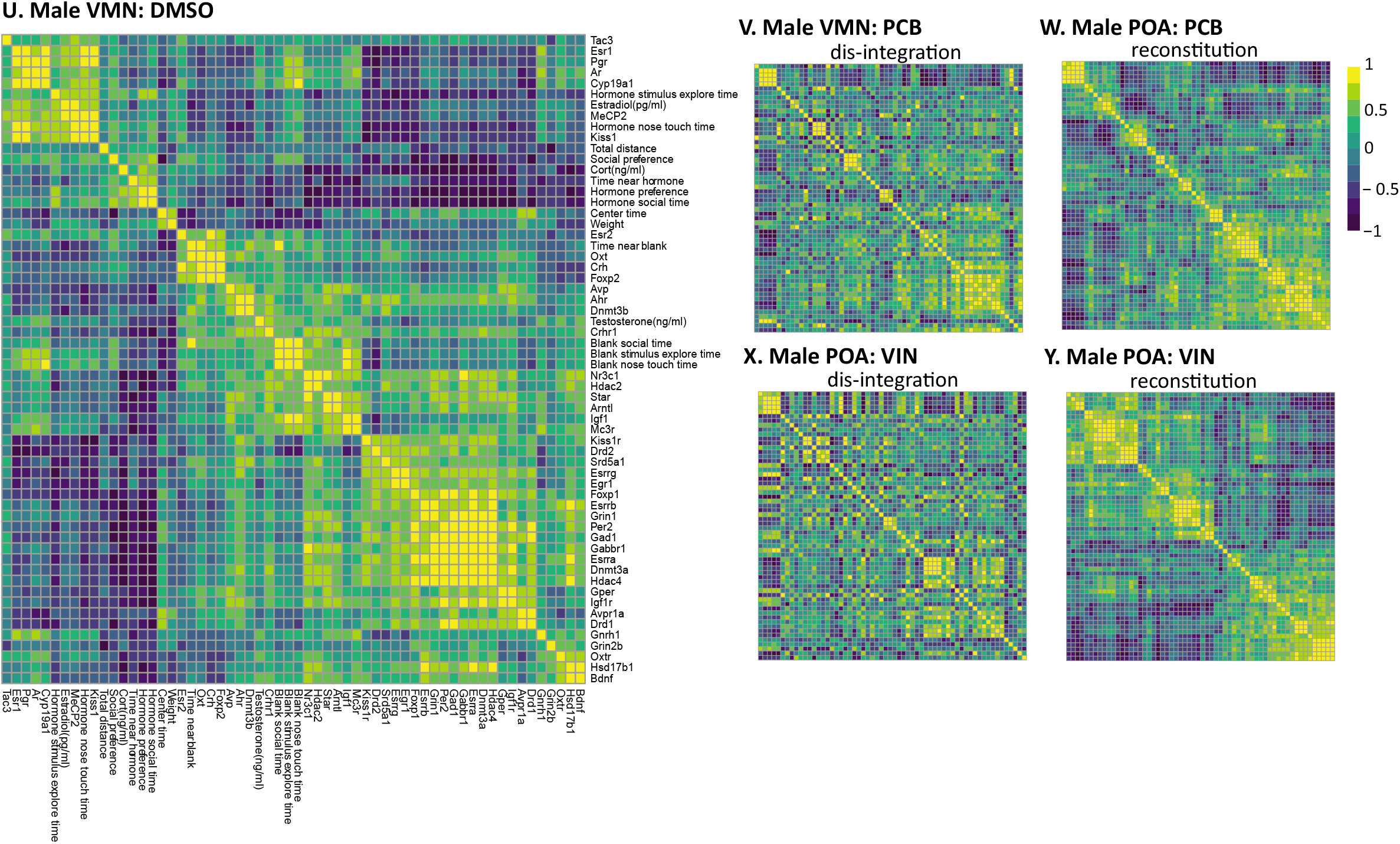
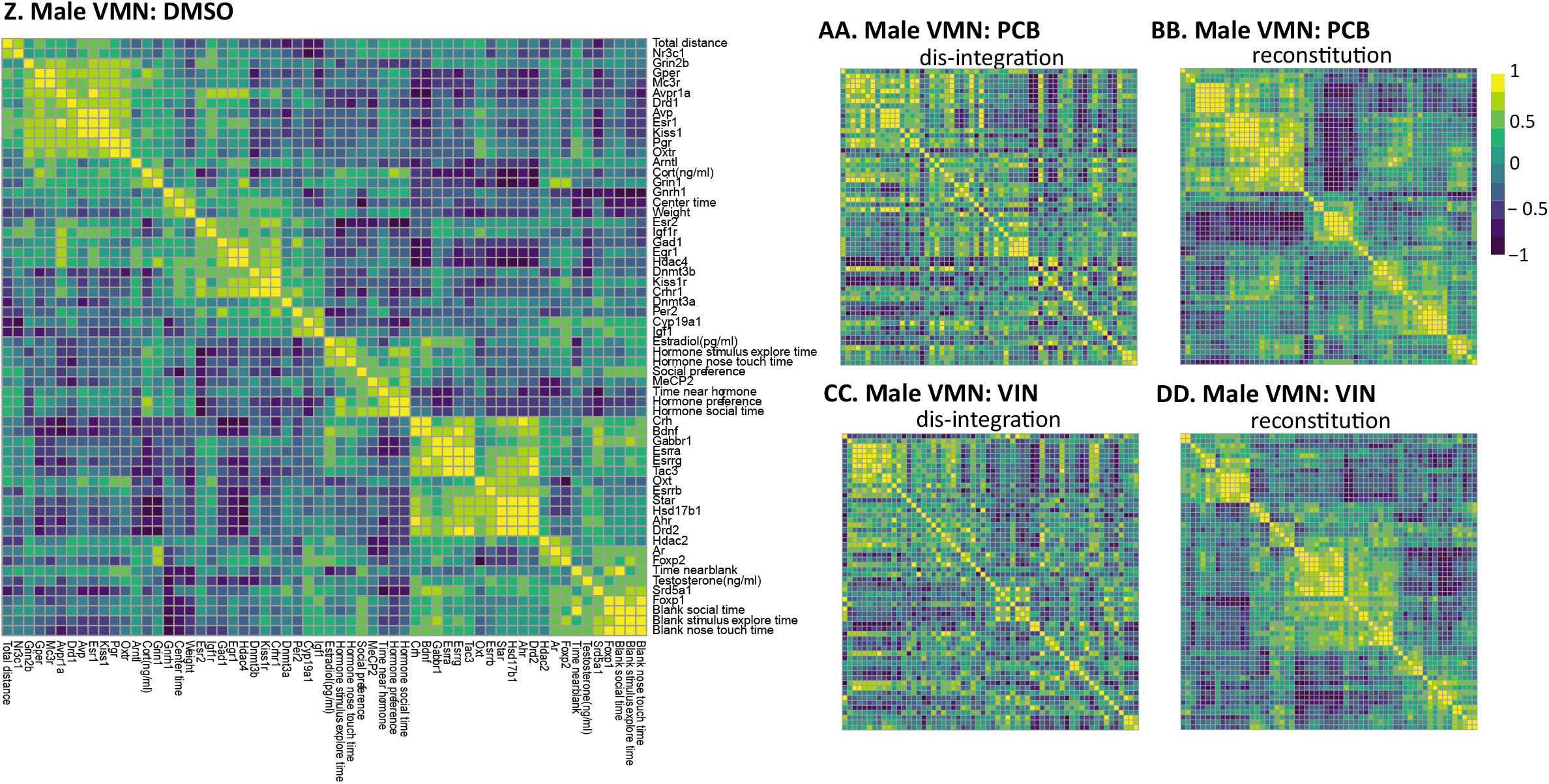

**Supplementary Figure 4.**
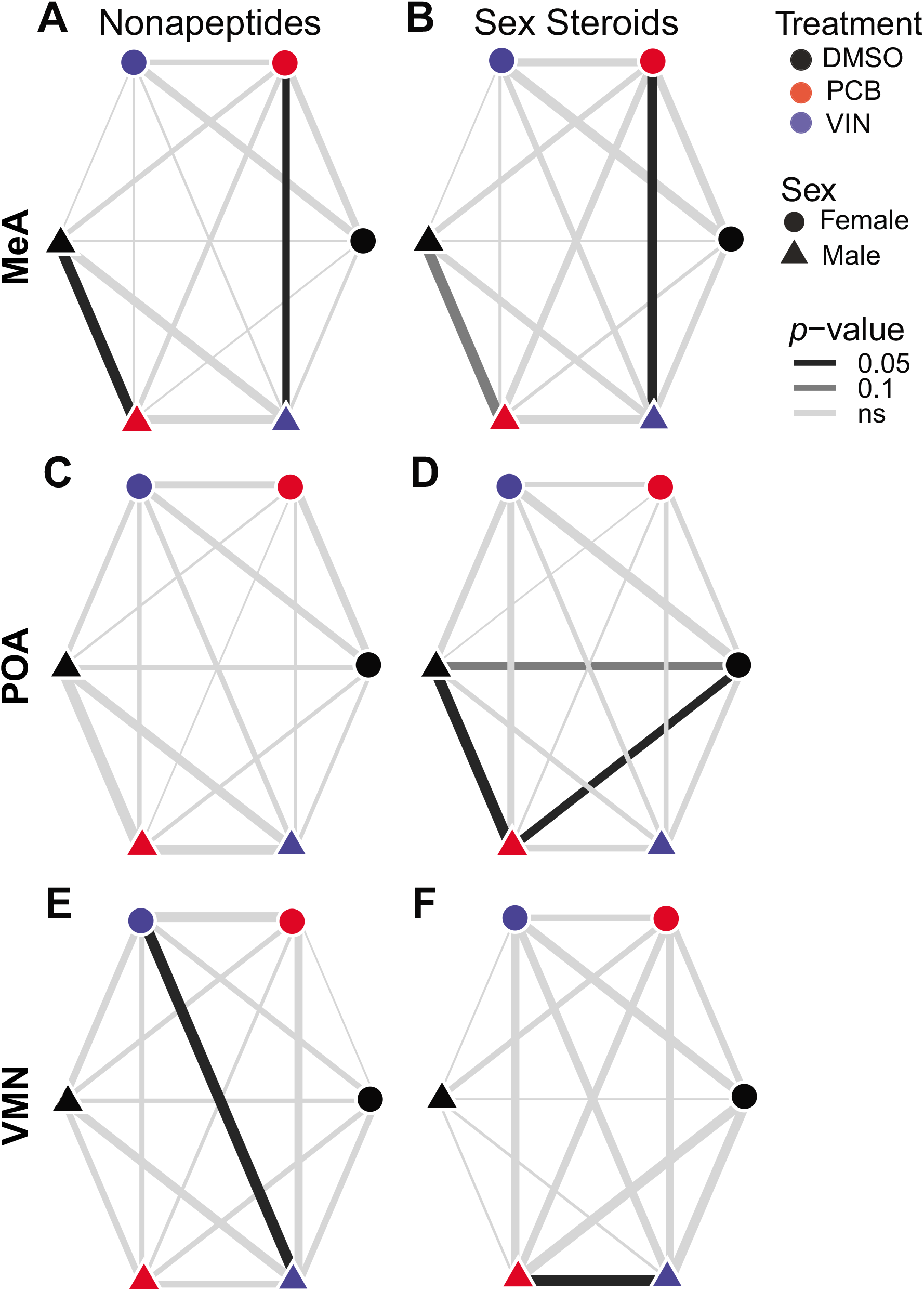

**Supplementary Table 1.**
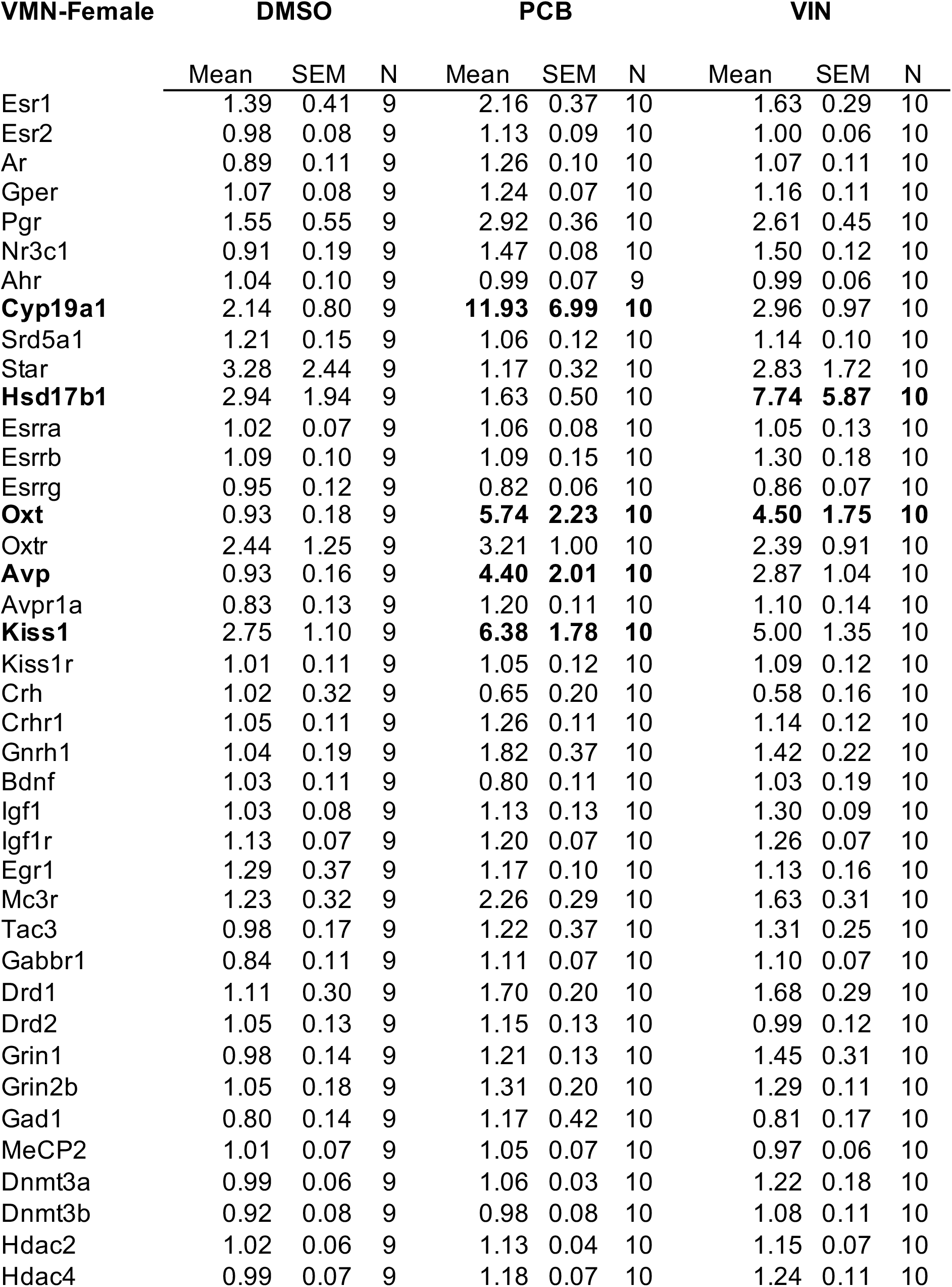
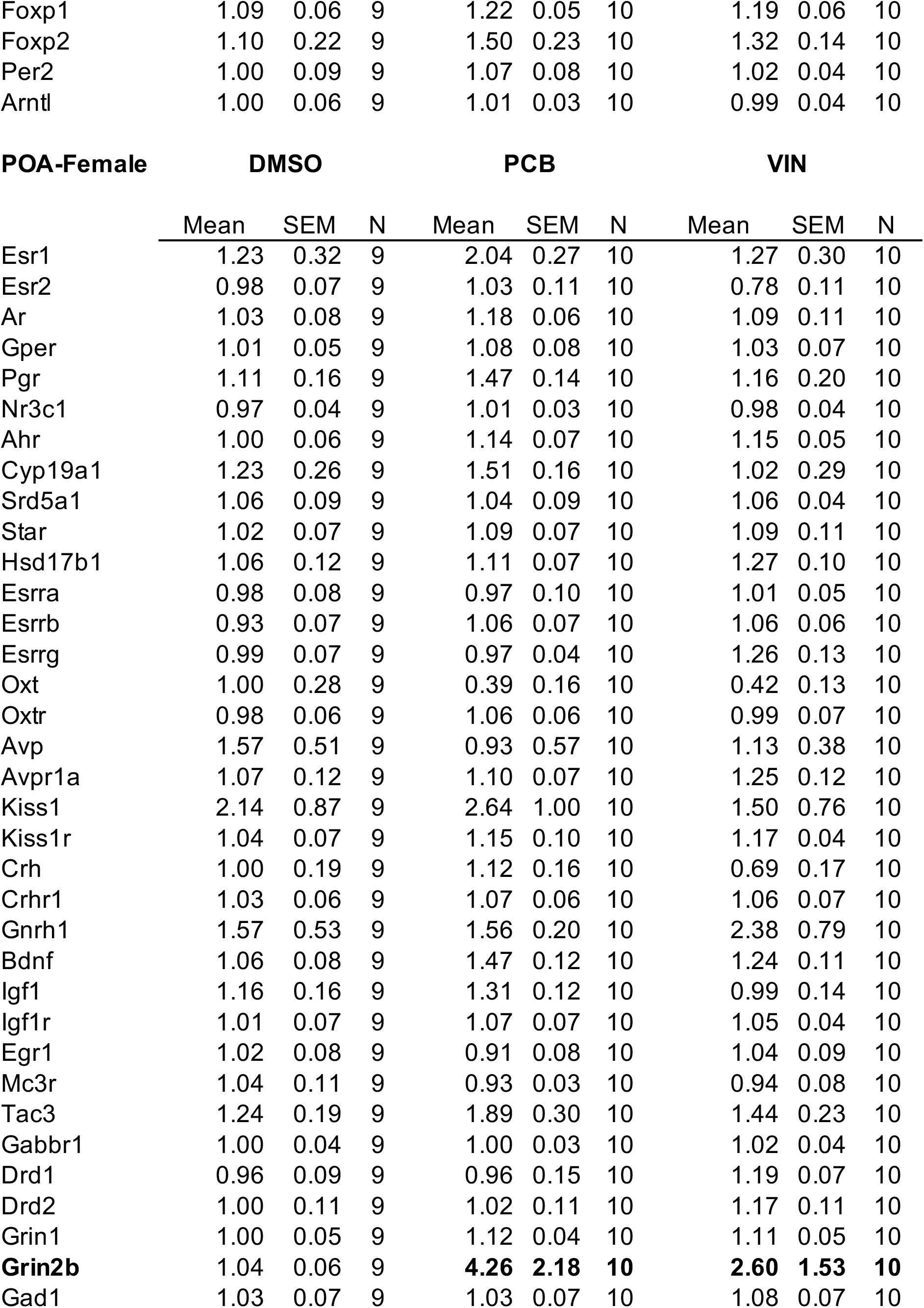
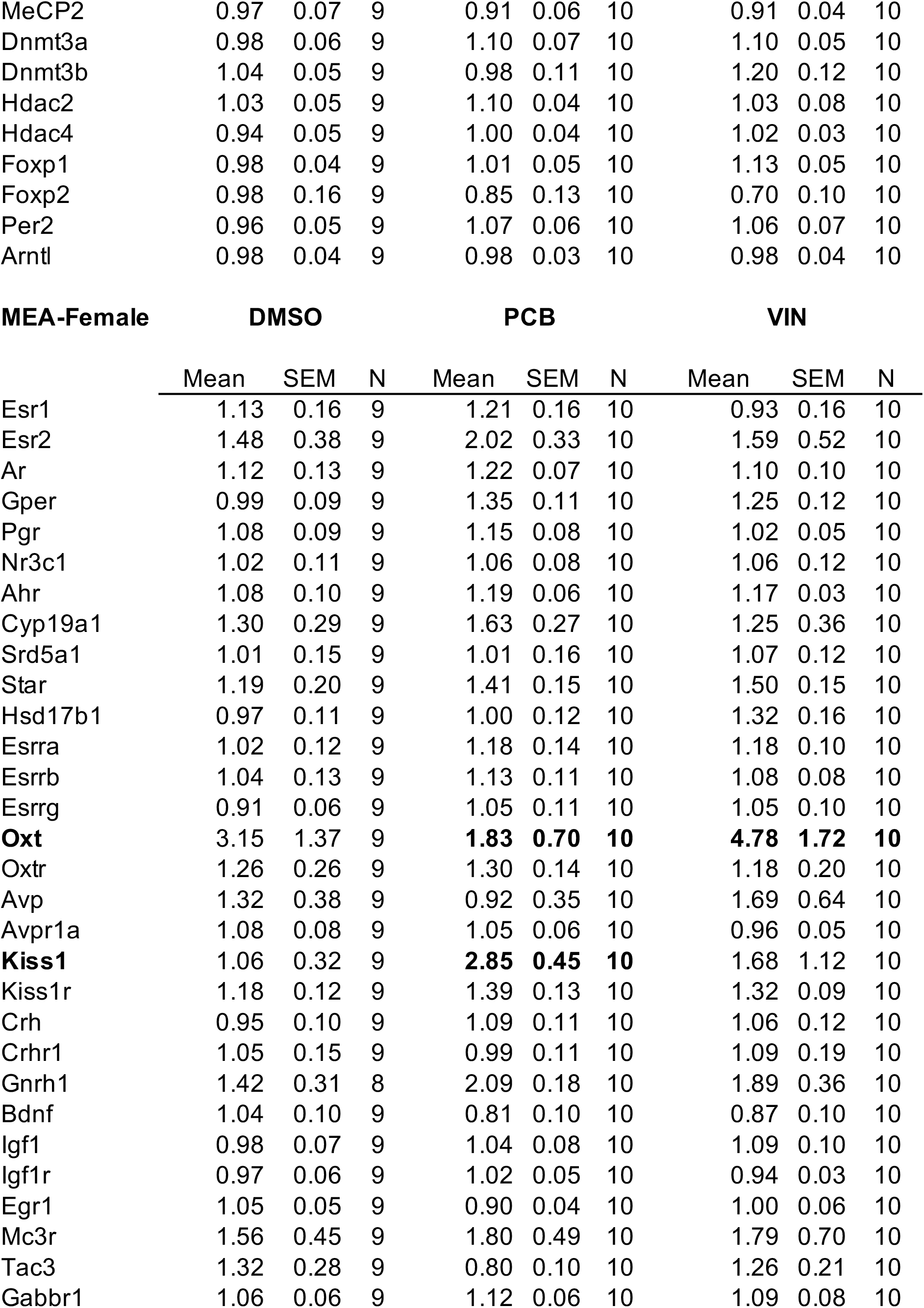
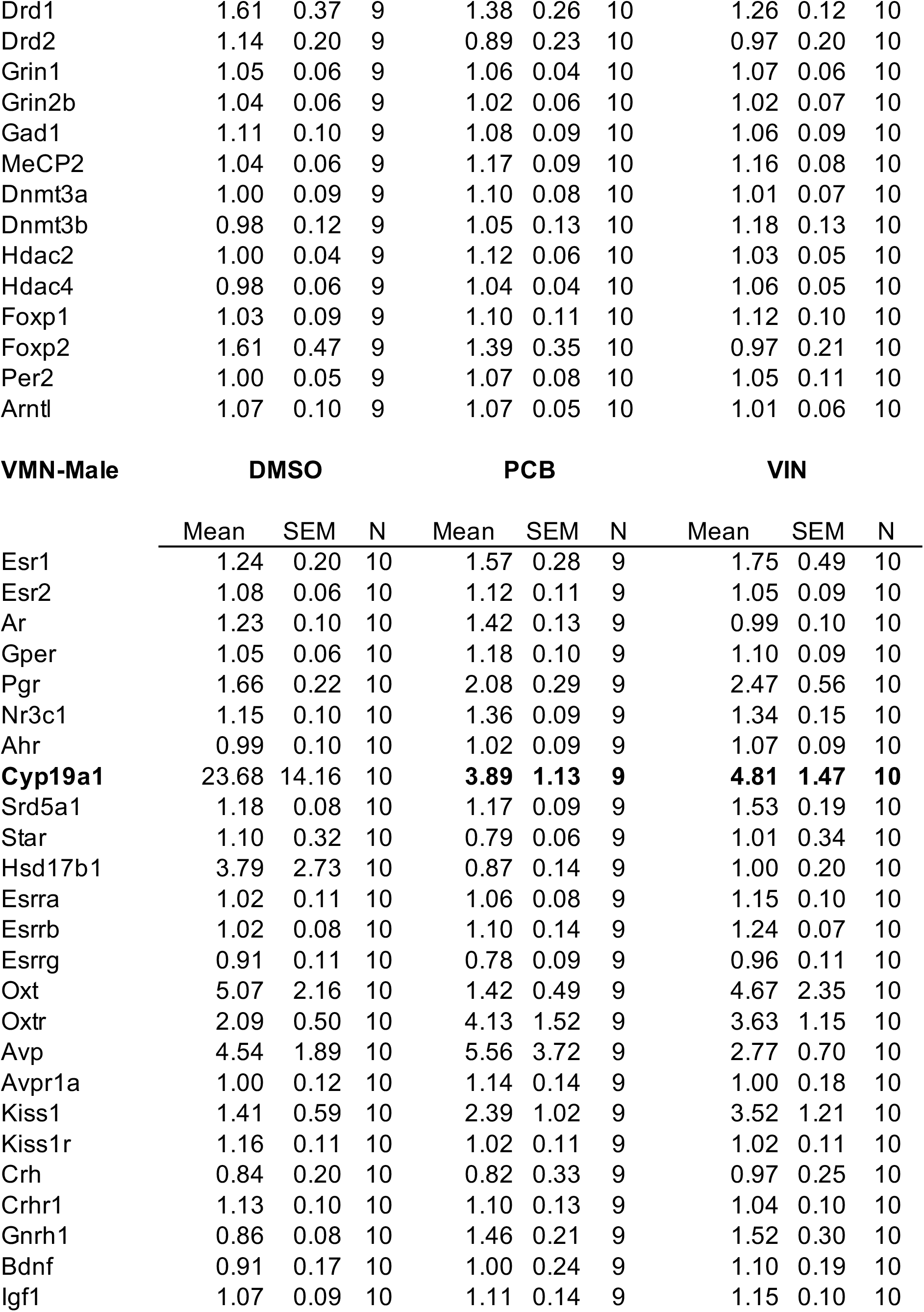
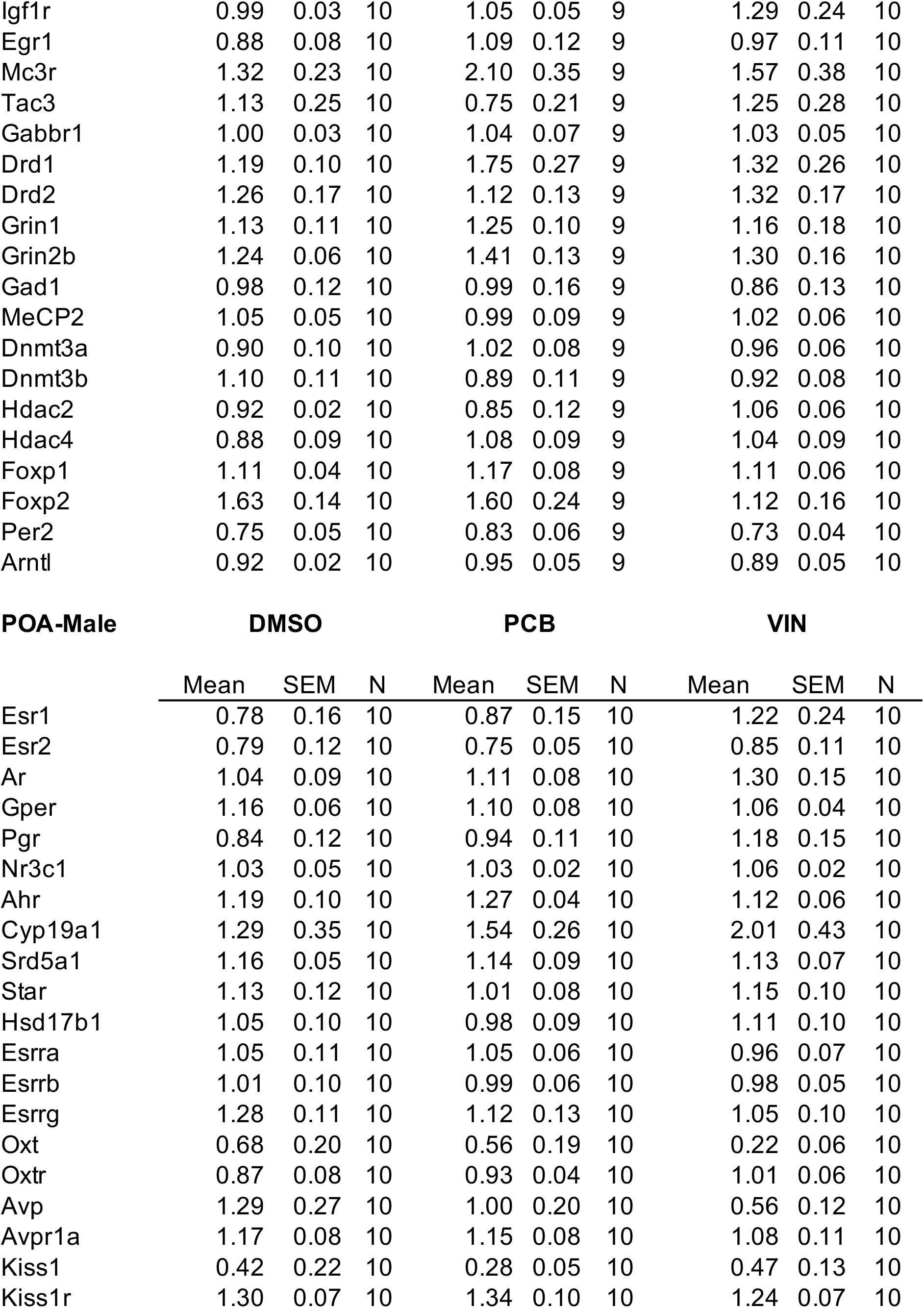
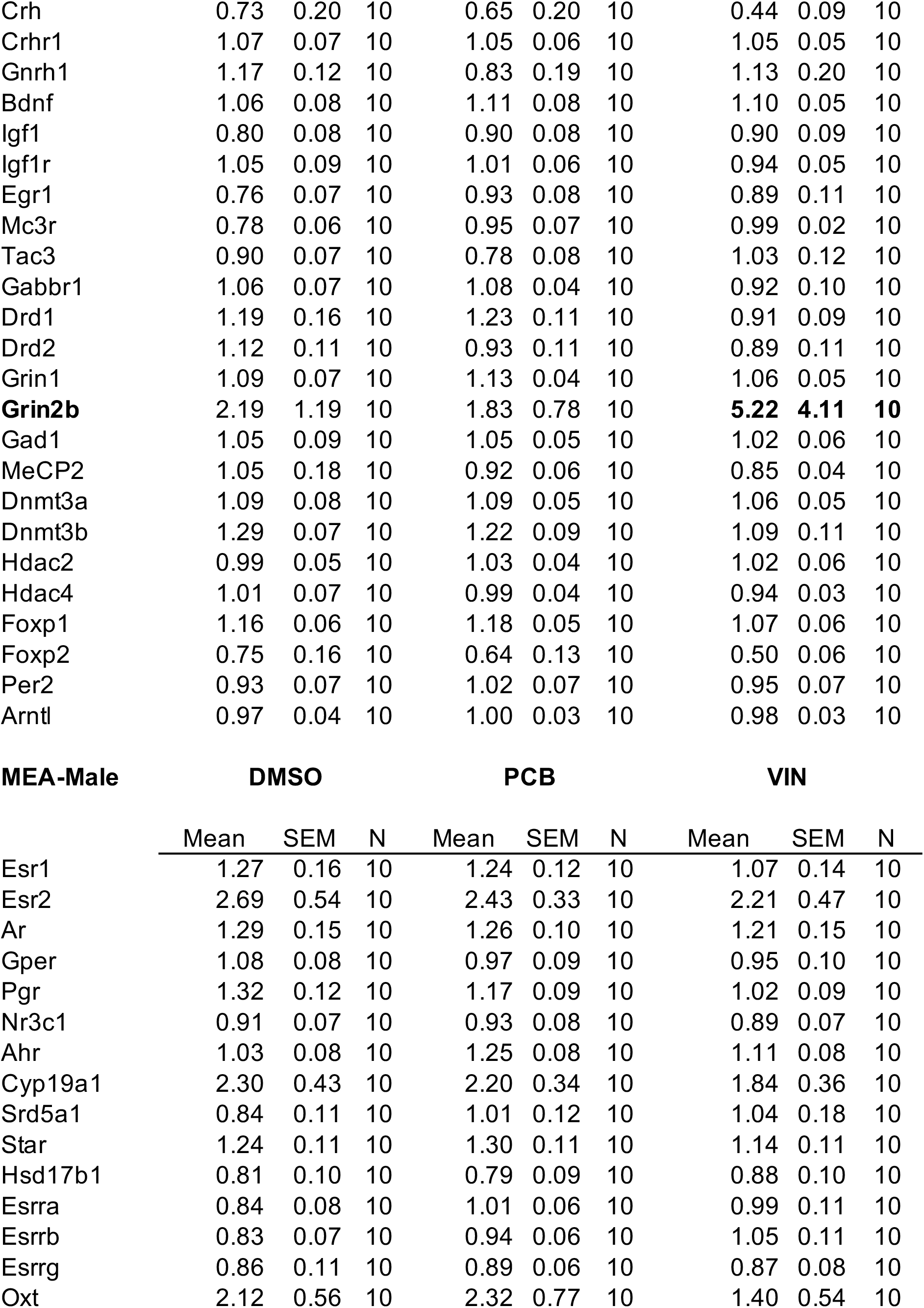
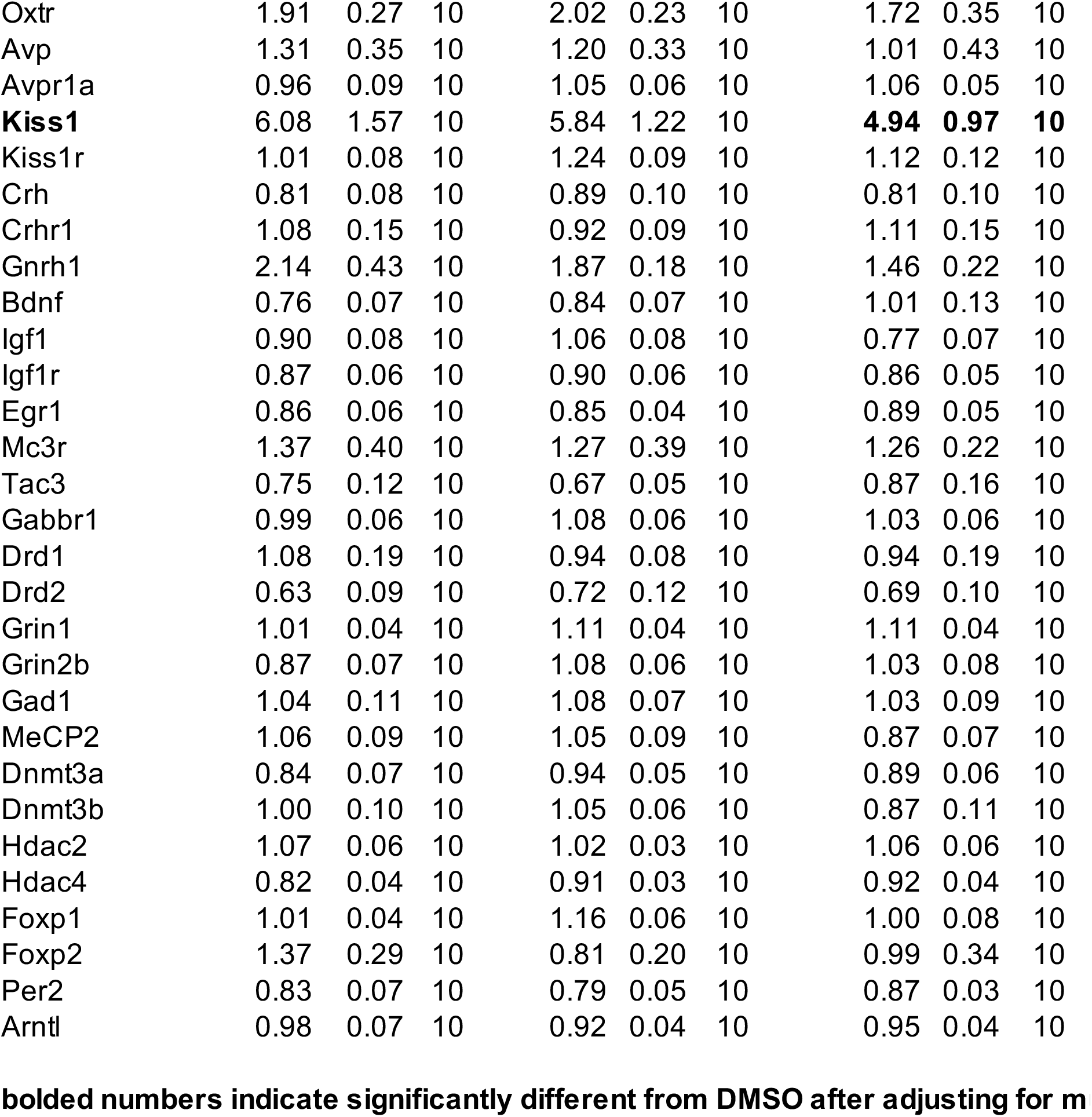
TLDA gene expression results

**Supplementary Table 2.**
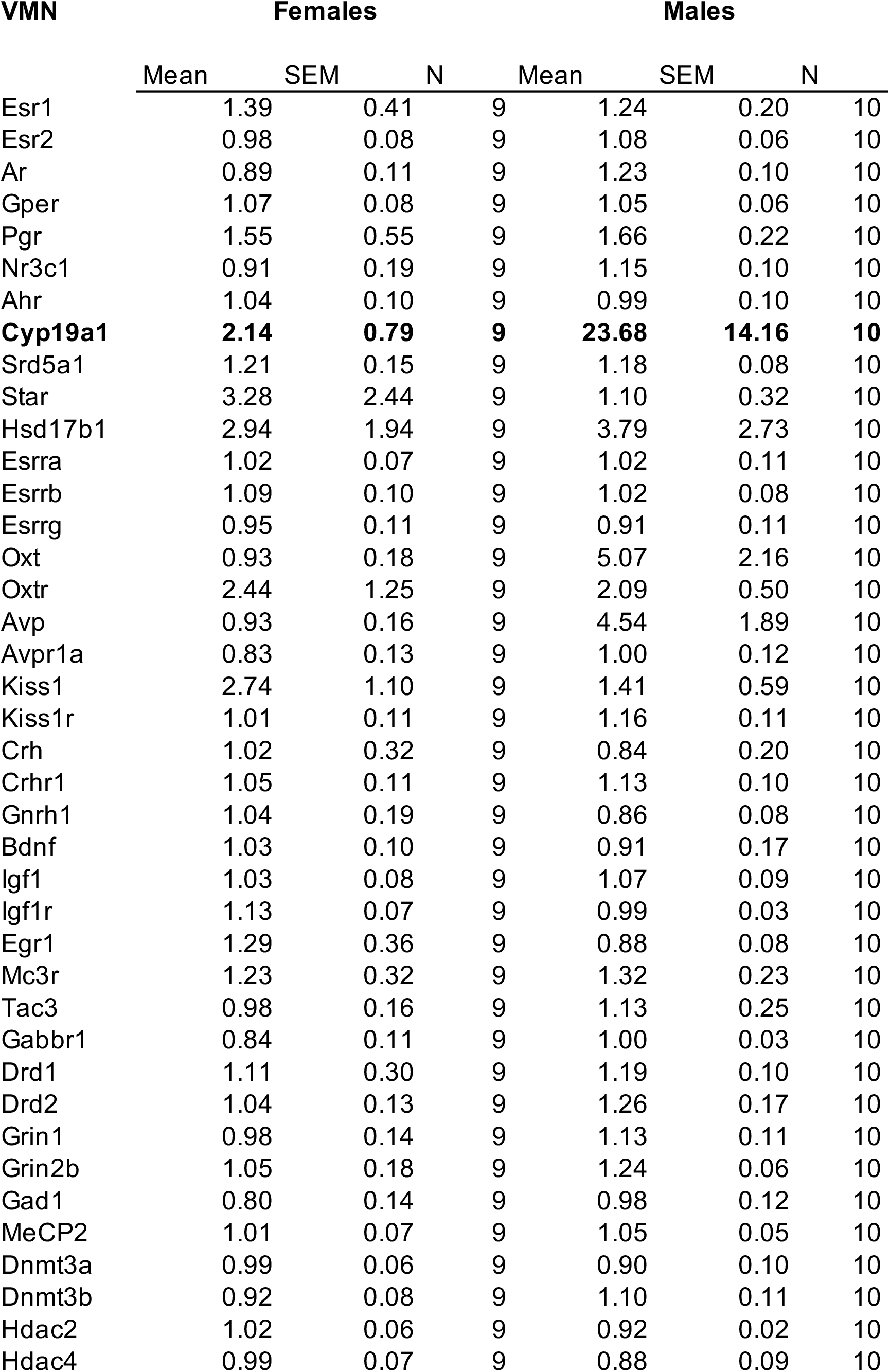
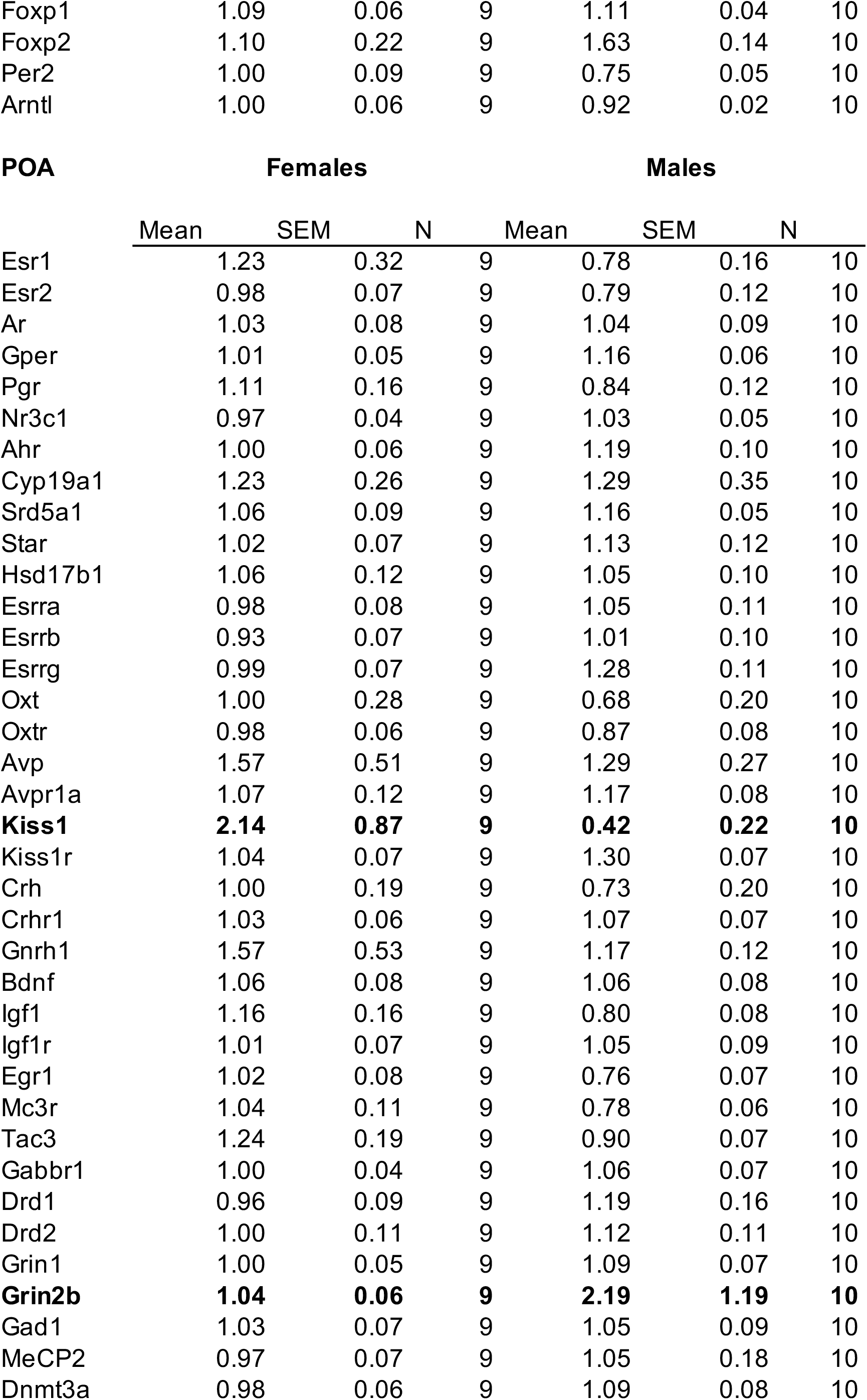
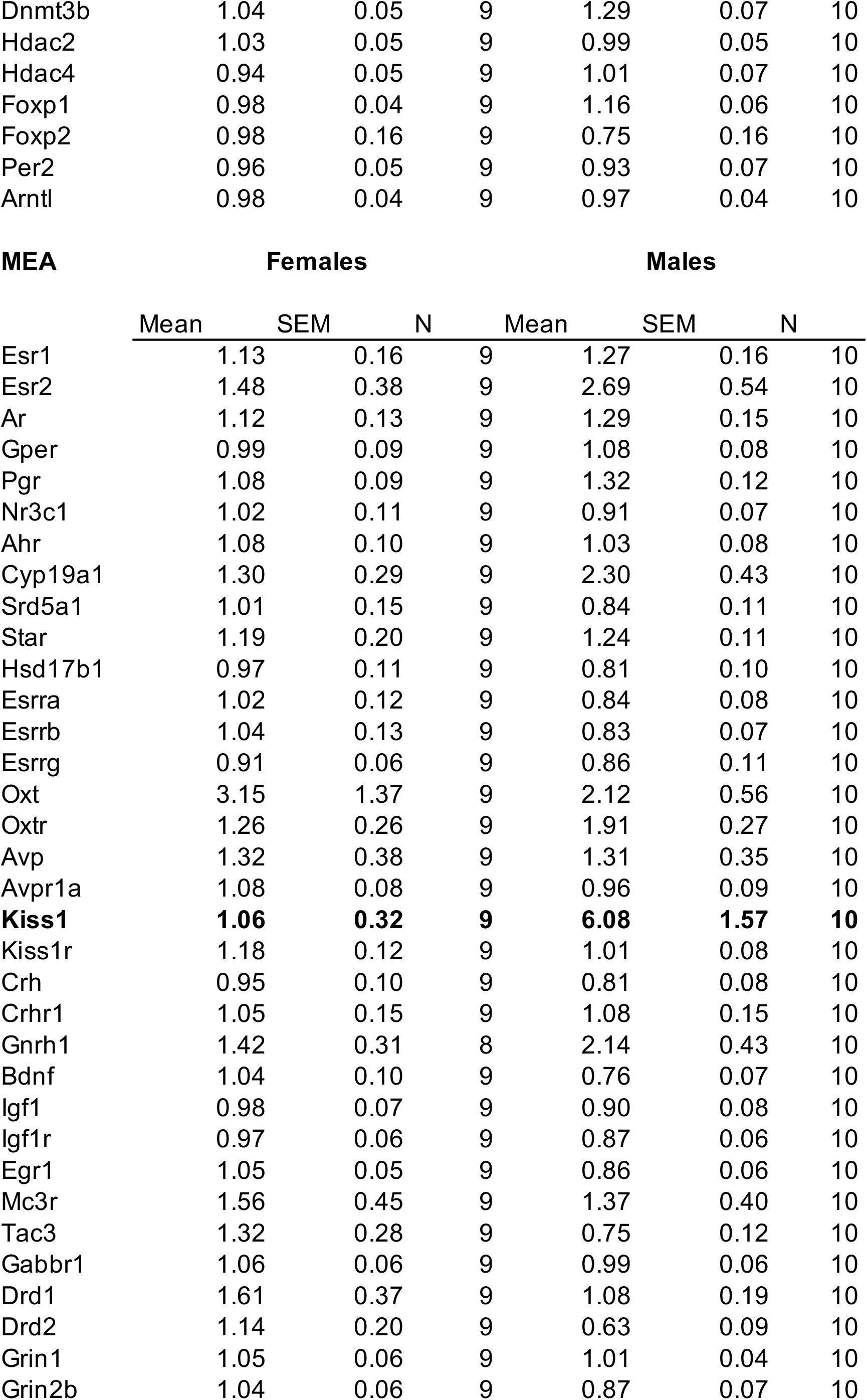
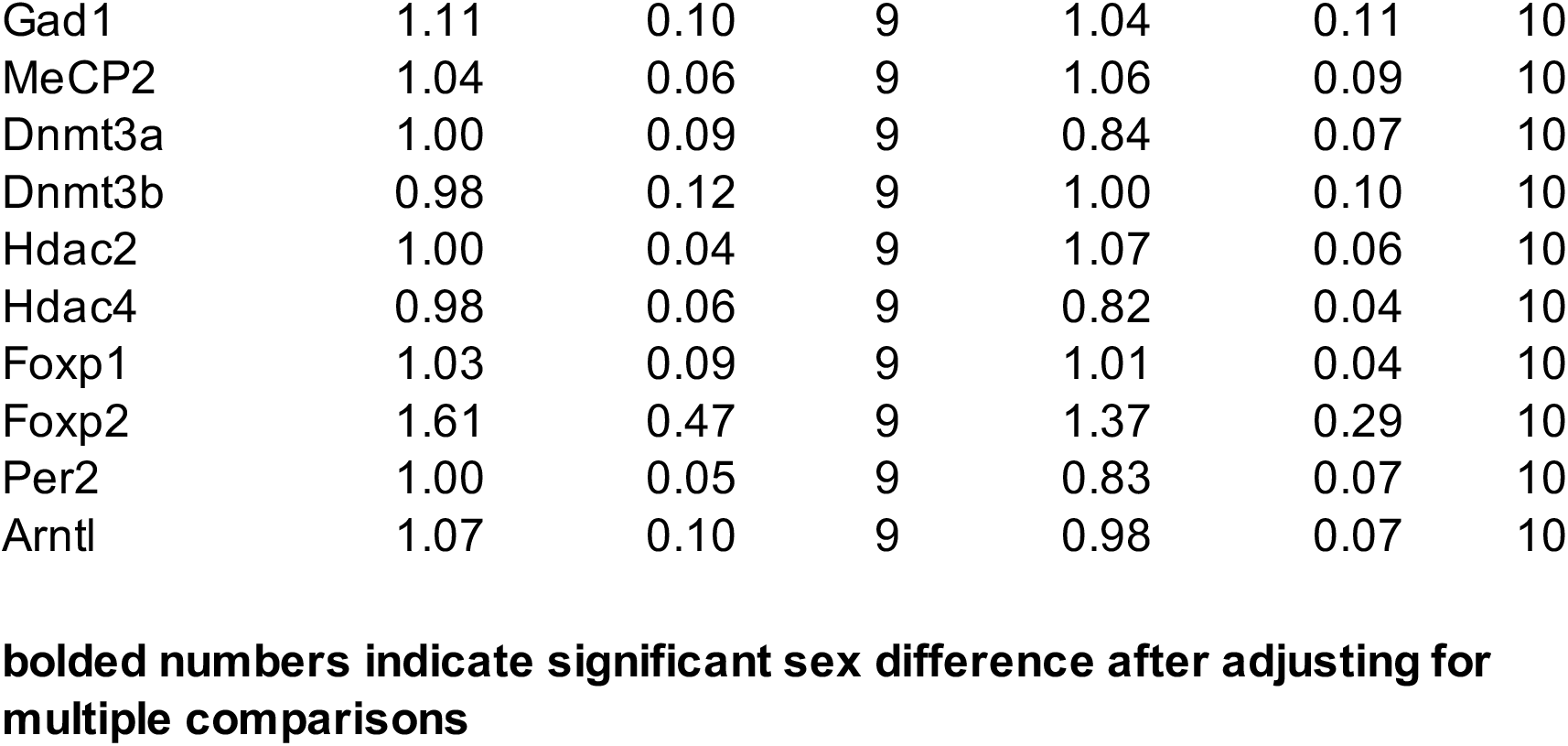
Sex differences in gene expression results

**Supplementary Table 3.**
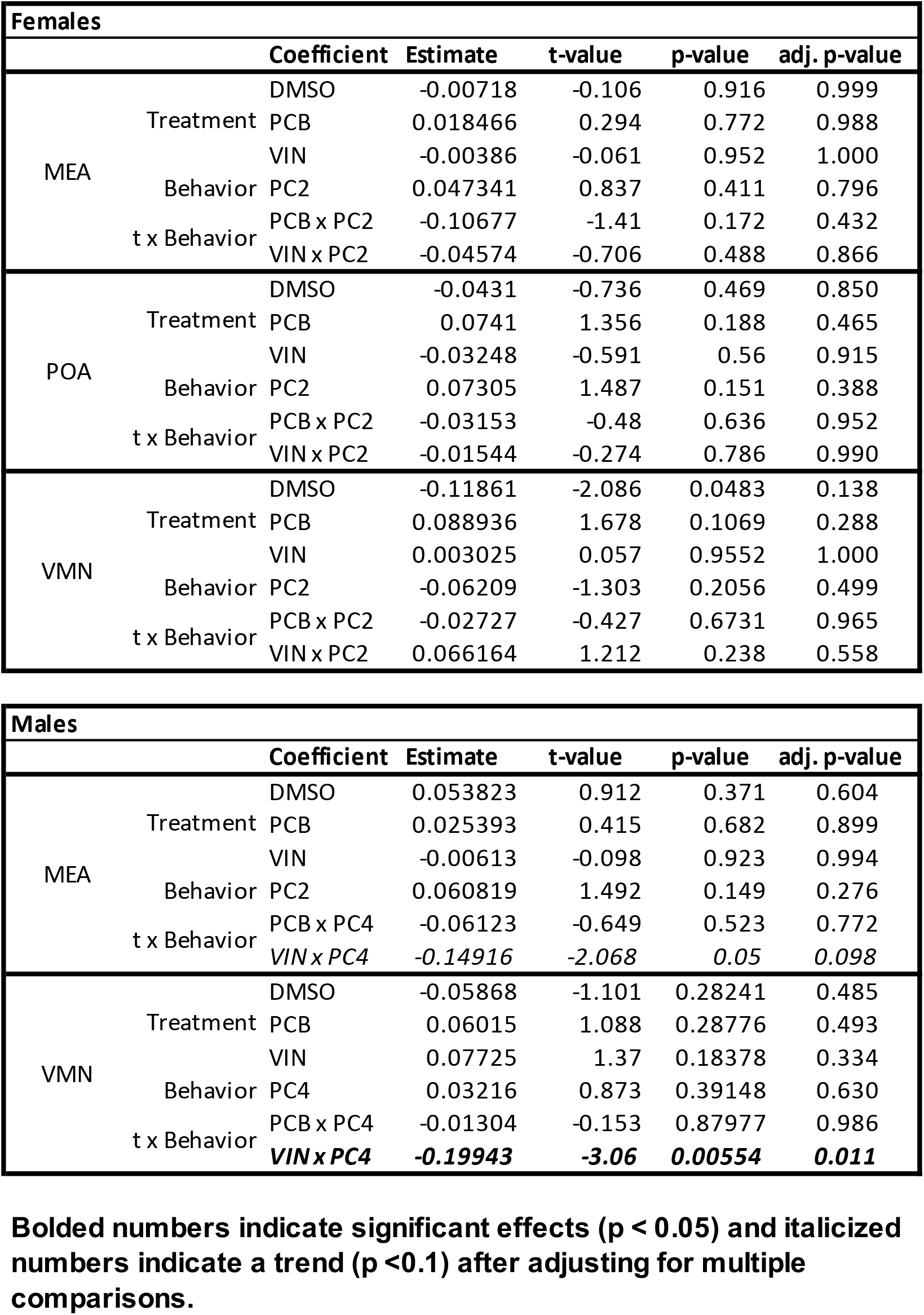
Gene co-expression, social interaction, and preference across treatments.

